# Classification and genetic targeting of cell types in the primary taste and premotor center of the *Drosophila* brain

**DOI:** 10.1101/2021.08.06.455433

**Authors:** Gabriella R. Sterne, Hideo Otsuna, Barry J. Dickson, Kristin Scott

**Affiliations:** University of California, Berkeley, United States; Janelia Research Campus, Howard Hughes Medical Institute, United States; Queensland Brain Institute; University of Queensland, St Lucia, Queensland, Australia

## Abstract

Neural circuits carry out complex computations that allow animals to evaluate food, select mates, move toward attractive stimuli, and move away from threats. In insects, the subesophageal zone (SEZ) is a brain region that receives gustatory, pheromonal, and mechanosensory inputs and contributes to the control of diverse behaviors, including feeding, grooming, and locomotion. Despite its importance in sensorimotor transformations, the study of SEZ circuits has been hindered by limited knowledge of the underlying diversity of SEZ neurons. Here, we generate a collection of split-GAL4 lines that provides precise genetic targeting of 138 different SEZ cell types in *D. melanogaster*, comprising approximately one third of all SEZ neurons. We characterize the single cell anatomy of these neurons and find that they cluster by morphology into six supergroups that organize the SEZ into discrete anatomical domains. We find that the majority of local SEZ interneurons are not classically polarized, suggesting rich local processing, whereas SEZ projection neurons tend to be classically polarized, conveying information to a limited number of higher brain regions. This study provides insight into the anatomical organization of the SEZ and generates resources that will facilitate further study of SEZ neurons and their contributions to sensory processing and behavior.

## Introduction

Elucidating the neural architecture that underlies sensorimotor transformations for behavior requires the ability to resolve and manipulate neural circuits with single cell precision. With a tractable number of neurons and well-developed genetic tools, *Drosophila melanogaster* is an excellent animal in which to investigate the basic principles of sensorimotor processing. Recent electron microscopy datasets provide unprecedented synaptic resolution of approximately 100,000 neurons that comprise the adult *D. melanogaster* brain (Scheffer et al., 2020; Zheng et al., 2018). If coupled with resources that provide genetic access to single neurons, this detailed anatomy may be probed to study complex circuits underlying sensory processing and behavior.

The subesophageal zone (SEZ) of the brain plays a critical role in many sensory driven behaviors. Defined as the brain tissue below the esophageal foramen, it is situated in a central location between the motor circuits of the ventral nerve cord (VNC) and the higher order brains regions of the supraesophageal zone (Ito et al., 2014). The SEZ participates in many context-dependent motor actions, including feeding, grooming, and locomotion, with evidence suggesting that it is involved in action selection (Bidaye et al., 2020; Flood et al., 2013; Gordon and Scott, 2009; Hampel et al., 2017, 2015; Mann et al., 2013; Manzo et al., 2012; Marella et al., 2006; Tastekin et al., 2015; Wang et al., 2004). It receives direct sensory input from axonal arbors of gustatory and mechanosensory peripheral neurons and indirect input from pheromone-sensing neurons (Ling et al., 2014; Thistle et al., 2012; Thorne et al., 2004; Toda et al., 2012; Wang et al., 2004; Weiss et al., 2011; Zhang et al., 2013). Two major outputs of the SEZ are descending neurons that convey information to the VNC and motor neurons that control the movement of the proboscis and antennae (McKellar et al., 2020; Namiki et al., 2018; Stocker et al., 1990). Recent work has delineated fascicle and neuropil-based columnar domains in the SEZ that are identifiable throughout development and has mapped sensory substructures in the SEZ (Hartenstein et al., 2018; Kendroud et al., 2018). Despite these advances, the cellular anatomy and function of SEZ neurons are largely unexplored.

Previous studies of SEZ cell types have relied on broad GAL4 lines or stochastic methods, which do not provide reliable access to individual neurons. Recent efforts using the split-GAL4 method in *Drosophila* have provided genetic access to libraries of single neurons in other brain regions, including the mushroom body, central complex, and lateral horn (Aso et al., 2014a; Dolan et al., 2019; Wolff et al., 2015). In this intersectional method, two different enhancers are used to independently drive expression of either the GAL4 transcriptional activation domain (AD) or DNA-binding domain (DBD). These domains heterodimerize through leucine zipper fragments and drive transgene expression restricted to the intersection of the two expression patterns (Luan et al., 2006). Thus, split-GAL4 reagents may be rationally designed if they are constructed using enhancers with known expression patterns. To systematically probe the cellular anatomy of the SEZ and to enable genetic dissection of SEZ neural circuits, we set out to create a library of genetic reagents to label individual SEZ cell types using the split-GAL4 method.

Here, we report the creation of 283 split-GAL4 lines that we collectively term the SEZ Split-GAL4 Collection. We estimate that this collection targets nearly one third of all neurons with cell bodies in the SEZ. Morphological clustering of the identified cell types reveals six layered (anterior to posterior) and stacked (inferior to superior) domains of organization in the SEZ. Furthermore, polarity analysis shows that many SEZ interneurons have inputs and outputs on the same processes, whereas SEZ projection neurons tend to be classically polarized, with inputs and outputs located in clearly distinct regions of the neuronal arbors. Taken together, the genetic reagents described here provide a valuable resource to investigate how diverse sensory inputs are processed by local SEZ circuitry to control specific behaviors.

## Results

### The SEZ contains about 1700 neurons

We set out to determine the number of neurons in the SEZ to inform the generation and assessment of split-GAL4 lines. The SEZ contains four anatomical subregions: the gnathal ganglia (GNG), saddle (SAD), antennal mechanosensory and motor center (AMMC), and the prow (Figure 1A and B), which, in general, are composed of cells from the tritocerebral, mandibular, maxillary, and labial neuromeres. To roughly estimate of the number of neurons in the SEZ, we assessed the number of neuronal cell bodies in each of these subesophageal neuromeres. This approach is likely to be an underestimate as this method does not account for deutocerebral contributions to SEZ neuron number (Boyan et al., 2003). We used a single-cell transcriptome atlas of the *D. melanogaster* brain (Davie et al., 2018) to determine the relative proportions of neurons expressing SEZ neuromere-specific markers. Then, we converted these proportions into neuron number estimates by counting cell bodies labeled by available Hox-gene drivers (Simpson, 2016) in individual *D. melanogaster* brains.

**Figure 1.**
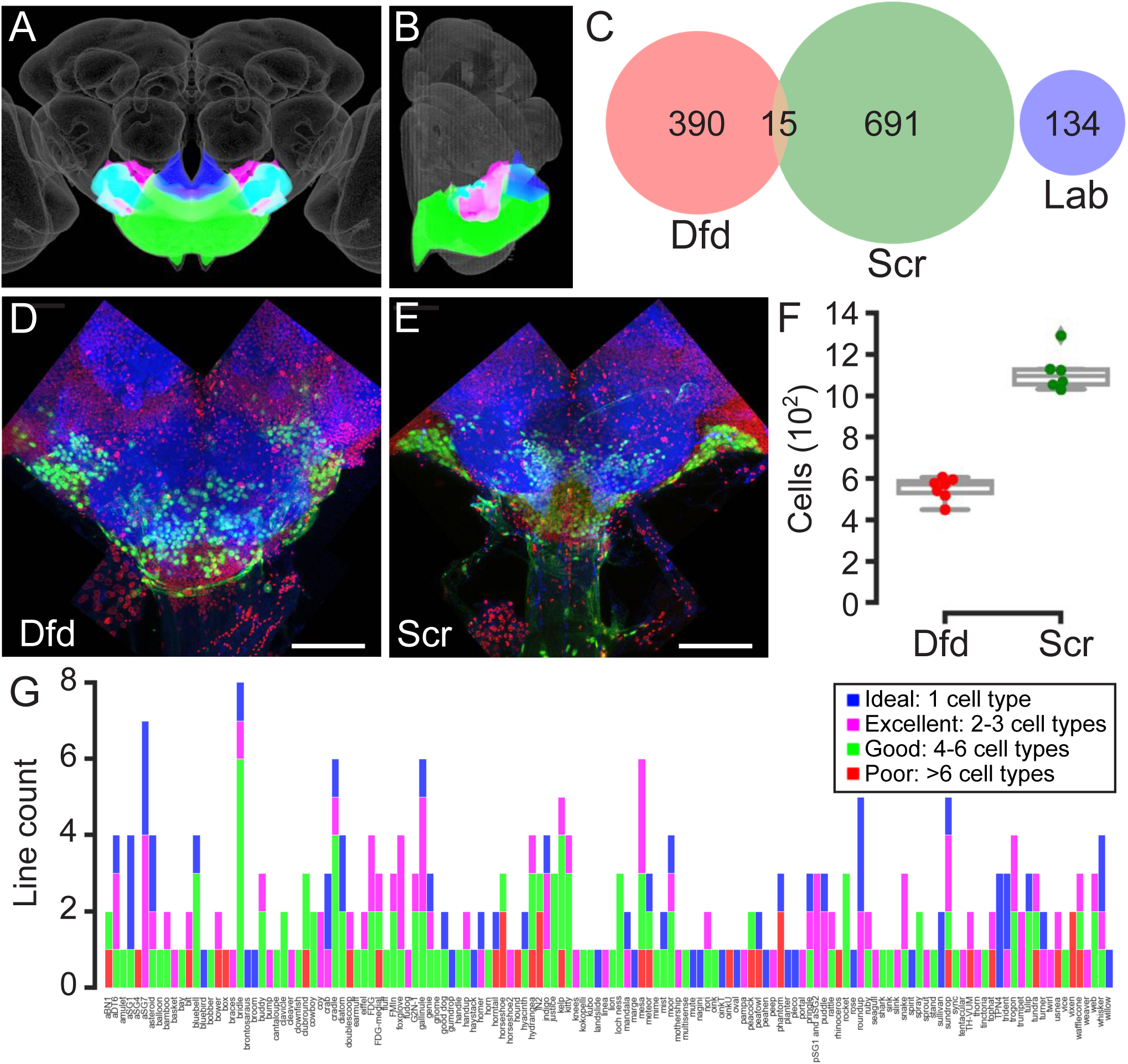
Coverage and quality of the SEZ Split-GAL4 Collection. (A, B) Anterior (A) and medial (B) views of the central brain of the *Drosophila melanogaster* adult showing the location of the gnathal ganglia (GNG, green), saddle (SAD, fuchsia), antennal mechanosensory and motor center (AMMC, cyan), and prow (royal blue) in relation to the JRC 2018 unisex brain template (grey). Together, the GNG, SAD, AMMC, and PRW compose the SEZ. (C) Venn diagram of single cells that with detectable *Dfd* (red), *Scr* (green), and/or *lab* (blue) as assessed with a single-cell transcriptome atlas. (D, E) Example stitched images used to count the number of cells expressing Dfd-LexA or Scr-LexA. LexAop-nls-GCAMP6s (green) driven by Dfd-LexA (D) or Scr-LexA (E) in the adult central brain. All nuclei are labeled with His2Av-mRFP (red) and neuropil is labeled with nc82 (blue). Scale bar 50 μm. (F) Box plots displaying counts of cell bodies labeled by both His2Av-RFP and LexAop-nls-GCaMP6s when driven with Dfd-LexA (n=7) or Scr-LexA (n=6). Whiskers denote spread of samples within 1.5 interquartile range from the mean. (G) The number of ideal (royal blue), excellent (fuchsia), good (green), and poor (red) split-GAL4 lines included in the collection by targeted neuron type.

We first examined the relative number of cells and neurons in each of the subesophageal neuromeres. We filtered single-cell RNA-sequencing data from the transcriptome atlas (Davie et al., 2018) to include only cells with detectable levels of any of three homeobox-containing transcription factors that are expressed in specific neuromeres: *Deformed* (*Dfd*) (mandibular and maxillary neuromeres), *Sex combs reduced* (*Scr*) (labial neuromere), or *labial* (*lab*) (tritocerebral neuromere) (Hirth et al., 1998; Kumar et al., 2015). Of the 56,902 high-quality cells represented in the atlas, 390 are *Dfd*-positive, 691 are *Scr*-positive, 134 are *lab*-positive, and 15 express both *Dfd* and *lab* (Figure 1C). Together, 1,230 cells in the atlas express these neuromere-specific markers, with 1,182/1,230 (96.1%) assigned to neuronal clusters based upon expression of neuronal genes. Although these results provide insight into relative numbers of cells in each subesophageal neuromere, they do not estimate neuron number in an individual brain because the single-cell RNA-seq atlas was constructed from multiple dissociated *D. melanogaster* brains.

To translate the proportions derived from single-cell RNA-sequencing data into an estimate of SEZ cell number, we directly counted cell nuclei in SEZ neuromeres in individual brains. Two knock-in LexA lines, Dfd-LexA and Scr-LexA (Simpson, 2016), were used to label cells in three of the four SEZ neuromeres, the mandibular and maxillary neuromeres and the labial neuromere, respectively (Figure 1D and E). We found that lab-GAL4, which is not a knock-in line, did not selectively label the tritocerebral neuromere, precluding cell counts of the fourth SEZ neuromere (Figure 1 – figure supplement 1). Adult female brains expressing nuclear localized GCaMP6s driven by either Dfd-LexA or Scr-LexA (neuromere-specific) and histone tagged with red fluorescent protein (RFP) under the control of the tubulin promoter (all cells) were used for visualization and machine-learning assisted quantification. Dfd-LexA labeled an average of 551 +/- 54 cells (n=7) while Scr-LexA labeled an average of 1115 +/- 94 cells (n=6) in the central brain (Figure 1F), generally consistent with the proportions seen in the transcriptome atlas. Using the direct counts of Dfd-LexA cells to estimate total SEZ cell number based on the proportions derived from single-cell RNA-sequencing, we would expect ∼1500-1850 cells in subesophageal neuromeres, ∼1450-1750 (96.1%) of which are likely to be neurons. Using our Scr-LexA counts to estimate total SEZ number, we would expect ∼1800-2100 cells, ∼1700-2000 of which are likely to be neurons. These estimates are roughly consistent with previous estimates of secondary SEZ neuron number based on neuroblasts, which predicted ∼2000 SEZ neurons (Kuert et al., 2014). We averaged the estimates based on Dfd and Scr counts to establish a final SEZ cell number estimate of ∼1800 cells, of which ∼1700 are neurons.

### The SEZ Split-GAL4 Collection provides genetic access to one third of all SEZ neurons

To characterize the morphology of individual SEZ cell types and to create a library of genetic reagents to provide specific access to these same cell types, we employed the split-GAL4 strategy (Luan et al., 2006). We used several strategies to identify novel SEZ cell types: 1) visual search through publicly available GAL4 collections (Jenett et al., 2012; Pfeiffer et al., 2008; Tirian and Dickson, 2017) 2) LexA-based MultiColor FlpOut (MCFO) single-cell labeling of Scr-LexA, and Dfd-LexA. 3) MCFO screening of subsets of the Rubin and Vienna Tile (VT) GAL4 collections with dense SEZ expression (Meissner et al., 2020; Nern et al., 2015), and 4) re-registration of images of individual SEZ cell types from mosaic analysis of broad GAL4 drivers, available on FlyCircuit (Chiang et al., 2011). Cell types innervating the AMMC were not included. Each novel cell type was given a unique (but not necessarily formulaic) name. Following cell type identification, we used the color depth maximum intensity projection (CDM) mask search tool (Otsuna et al., 2018) to select available split-halves that potentially labeled each cell type. After gathering a list of available split-halves likely to label a given cell type, we crossed all possible combinations of candidate ADs and DBDs and screened for split-GAL4 lines that specifically labelled the cell type of interest.

We screened ∼3400 split-GAL4 combinations using this strategy, which yielded 283 lines that provide precise access to single cell-types in the SEZ. The expression of each line is annotated to indicate the cell type that it was designed to target and the quality of the line (supplemental table). These split-GAL4 lines label 138 SEZ cell types, 129 of which have not been previously reported. The quality of each line was rated as ideal (labeling only a single SEZ cell class and no other neurons in the brain or VNC), excellent (labeling the cell type of interest and 1-2 other cell-types), good (labeling the cell type of interest and 3-5 other cell-types), or poor (labeling the cell type of interest plus more than 5 other cell types). Amongst the 283 split-GAL4 lines that were generated, 63 are ideal, 87 are excellent, 102 are good, and 31 are poor. Poor lines are included in the collection if they improve genetic access to the target cell type as compared to existing GAL4 lines. Each target cell type is covered by at least one split-GAL4 line. The number and quality of split-GAL4 lines per targeted cell type is shown (Figure 1G).

To evaluate the completeness of coverage achieved by the SEZ Split-GAL4 Collection, we compared the total number of neuronal cell bodies covered by the split-GAL4 lines with SEZ neuron number estimates. SEZ cell types fall into either unique or population classifications, where unique neurons encompass a single pair of cell bodies while population neurons are small groups of cell bodies with nearly identical arbors (Namiki et al., 2018). Therefore, one cell type may contribute one or multiple cell bodies per hemisphere. Taking this into consideration, the collection labels 510 neurons out of 1,700 estimated, arguing that the SEZ Split-GAL4 Collection provides approximately 30% coverage of all SEZ neurons. In addition, 17 split-GAL4 lines specifically target SEZ motor neurons of the proboscis, totaling 36 cell bodies (McKellar et al., 2020). Moreover, the Descending Interneuron (DN) Split-GAL4 Collection contains 41 DN cell types that comprise 242 additional cell bodies in the SEZ (out of 360 total DN cell bodies in the SEZ) (Namiki et al., 2018). Together, the SEZ Split-GAL4 Collection, the proboscis motor neuron split-GAL4s, and the DN Collection provide precise access to 46% of SEZ neurons (788/1,700). In summary, the SEZ Split-GAL4 Collection greatly improves genetic access to SEZ cell types, especially non-DN SEZ cell types. These split-GAL4 lines represent a substantial expansion of the knowledge of SEZ cell types and enable precise manipulation of the targeted cell types for behavioral, functional imaging, and morphological analyses. Confocal images of each line and instructions for requesting lines from the SEZ Split-GAL4 Collection can be found here: https://splitgal4.janelia.org/.

### Clustering of SEZ cell types reveals six cellular domains

To investigate SEZ organization at a cellular level, we used the NBLAST algorithm to perform automated clustering of SEZ cell types to define cell type supergroups (Costa et al., 2016). NBLAST computes a pairwise neuronal similarity score by considering the position and local geometry of a query and target neuron. By comparing neurons with NBLAST in an all-by-all matrix, we clustered them into morphologically similar groups to reveal SEZ substructure. To prepare neuron imagery for the NBLAST algorithm, a single, unilateral example of each cell type was imaged at high resolution using MCFO and registered to a common unbiased template (Bogovic et al., 2020; Nern et al., 2015). Each cell type example was then segmented, skeletonized, and presented on the right side of the brain. In total, 121 of the 138 SEZ cell types targeted by the collection are represented in this dataset. The remaining cell types were excluded from NBLAST analysis because MCFO images were not available. The expression pattern of the best split-GAL4 line targeting each cell type excluded from NBLAST analysis is shown (Figure 2 – figure supplement 1). After preprocessing, we computed an all-by-all similarity matrix for the represented cell types with NBLAST and hierarchically clustered the resulting NBLAST scores using Ward’s method (Costa et al., 2016) (Figure 2B). Ward’s method is an agglomerative hierarchical clustering method that groups items into clusters that minimize within-cluster variance. Ward’s joining cost, which is based on the variance of the data within a cluster, should increase significantly when distinct groups within the data are forced to join (Braun et al., 2010). Since the expected number of groups was not known beforehand, we analyzed Ward’s joining cost and the differential of Ward’s joining cost to quantitatively determine group number. We chose six groups due to the low joining cost and the increase in the differential of Ward’s joining cost when moving from six to five groups (Figure 2 – figure supplement 2).

**Figure 2.**
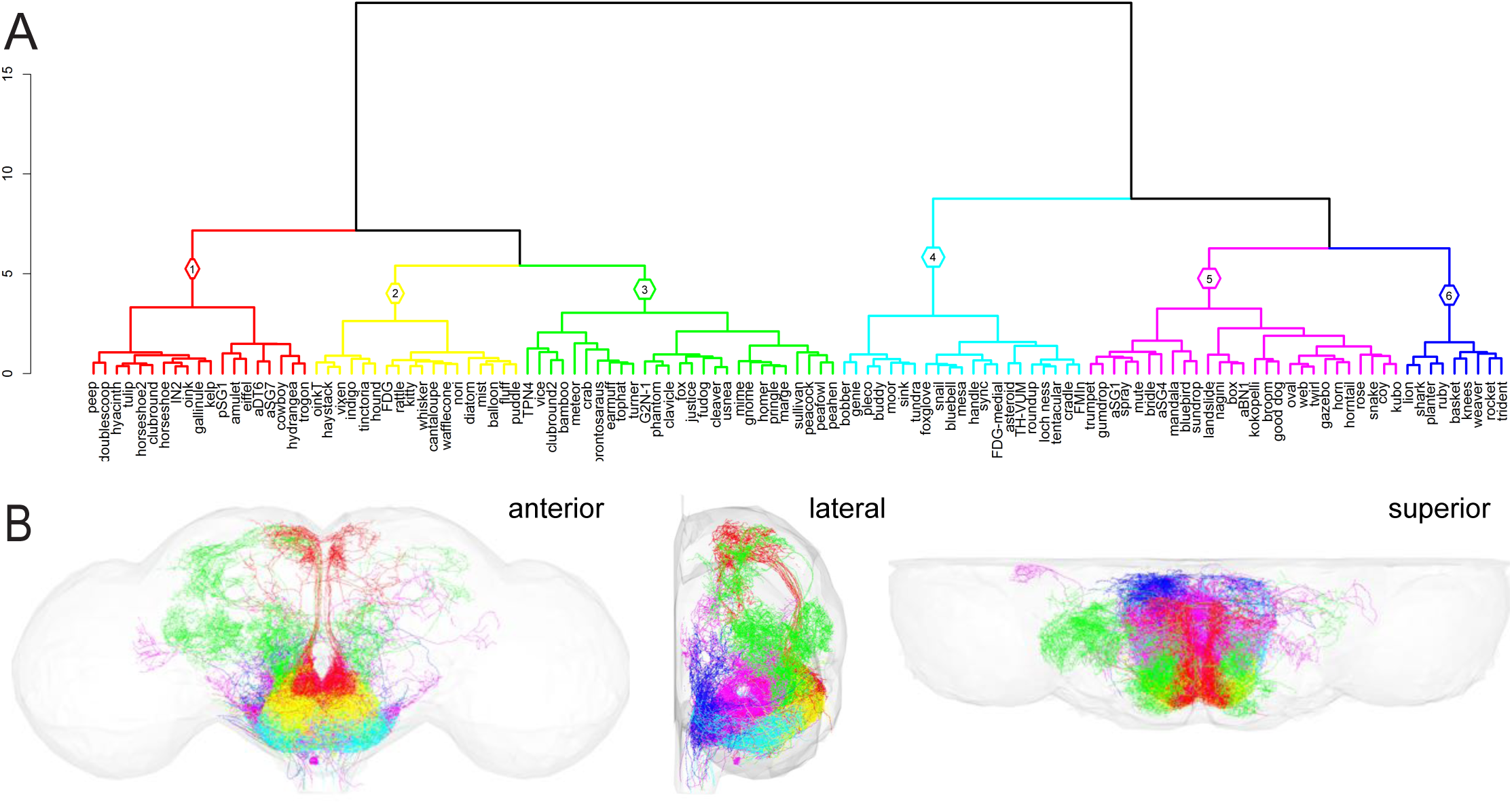
Hierarchical clustering of SEZ neuronal cell types (A) Clustering of SEZ neuron types with NBLAST reveals six distinct morphological groups: Group 1, red; Group 2, yellow; Group 3, green; Group 4, cyan; Group 5, fuchsia; Group 6, royal blue. Group number is denoted in the diamond above each cluster. (B) Morphology of all neuron types in each cluster plotted according to the color code in (A). Central brain neuropil (grey) is plotted for reference. Anterior (left), lateral (middle), and superior (right) views are shown.

The resulting supergroups share anatomical similarities and coordinates that reveal that the SEZ is organized into layered and stacked domains. Five of the six supergroups are layered from anterior to posterior: 1 and 2 most anterior, followed by 3, 5, and finally 6 most posterior. Groups 1 and 2 are in a similar anterior plane but Group 1 is positioned superior to Group 2. Group 4 sits below these domains, wrapping the inferior surface of the SEZ. A lateral view illustrates that Group 5 appears to form a “roll” shape and is surrounded by Group 3 anterior, Group 4 inferior, and Group 6 posterior. Cell types within the supergroups share morphological commonalities that may reflect functional similarities. We speculate about potential behavioral functions for each supergroup but acknowledge that the roles of the neurons described here are likely more diverse. For each group we show the morphology of an individual, segmented neuron for each cell type (Figures 3-8) as well as the pattern of the best split-GAL4 line for that cell type (Figures 3-8, figure supplement 1).

**Figure 3.**
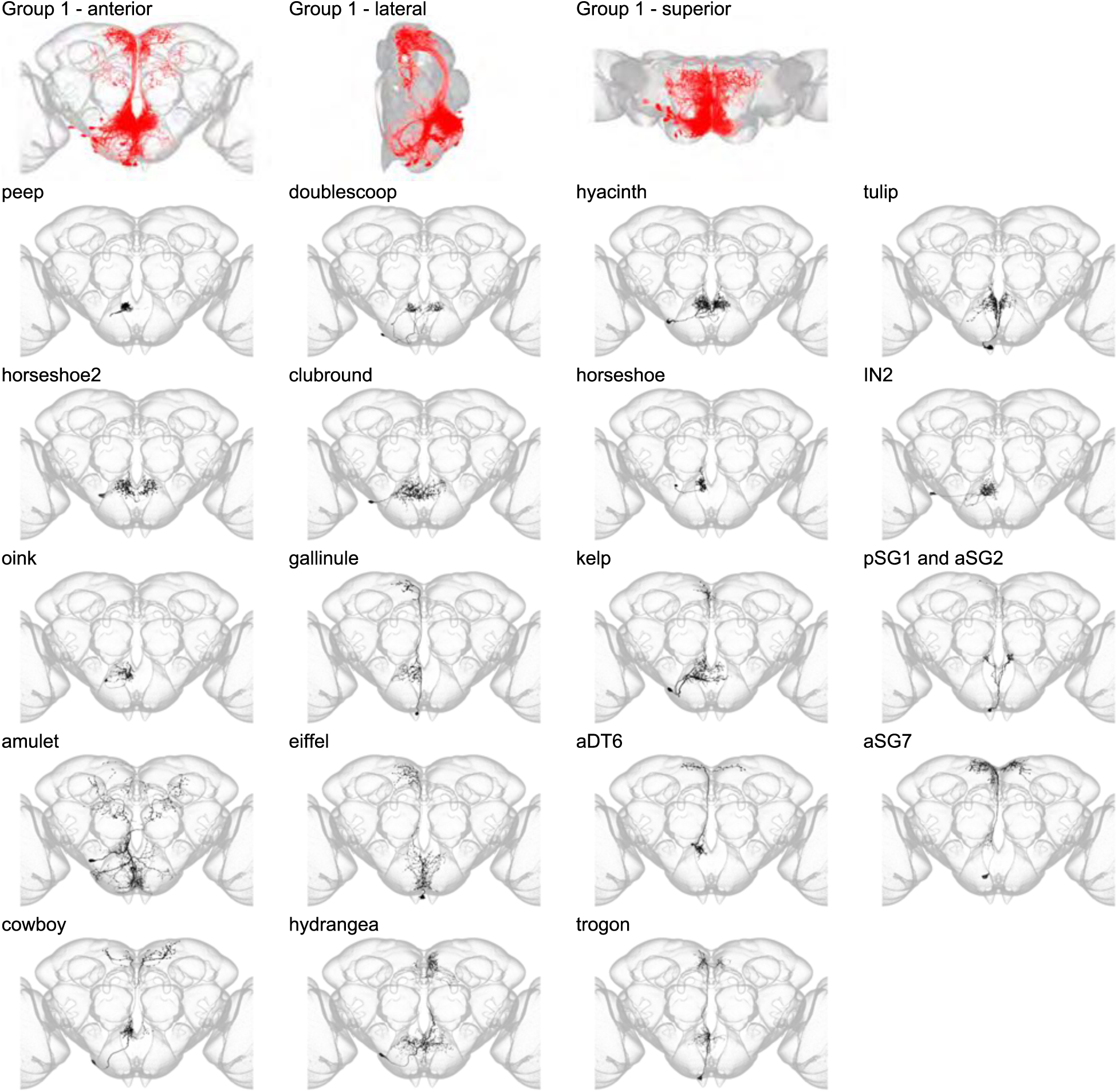
Morphology of neuron types in Group 1. Segmented example images for each neuron type in Group 1. The top row shows the morphology of all neuron types in Group 1 (red) overlaid in the 2018U coordinate space (grey) in anterior, lateral, and superior views. Below, the morphology of individual group members are shown separately. Individual neuron morphology is shown in black while the outline of the JRC 2018 unisex template is shown in grey. In Figures 3-8 the segmented neurons were imaged with a 63x objective and registered to the full-size JRC 2018 unisex template. The optic lobes have been partially cropped out of each panel.

Group 1 is composed of neurons that arborize in the prow and flange (Figure 3 and Figure 3 – figure supplement 1), the superior, anterior, and medial region of the SEZ. Based on their anatomical position, the 19 cell types that make up Group 1 may originate from the tritocerebral neuromere. Group 1 is neatly split into interneurons (first half of the group) and projection neurons (second half of the group). Notably, all projection neurons in Group 1 send their axons to the superior medial protocerebrum (SMP) (Figure 9). This group includes three previously morphologically described fruitless positive (Fru+) neuronal cell types: aSG2, aSG7, and aDT6 (Liu, Tianxiao, 2012; Yu et al., 2010). Because Group 1 projection neurons innervate the region of the SMP surrounding the pars intercerebralis (PI), we speculate that Group 1 neurons may impinge on neurosecretory neurons or function in energy and fluid homeostasis circuits. The proximity of Group 1 interneurons to previously described Interoceptive SEZ Neurons (ISNs) (Jourjine et al., 2016) and Ingestion Neurons (IN1) (Yapici et al., 2016) supports this hypothesis.

Group 2 contains only SEZ interneurons (Figure 4 and Figure 4 –figure supplement 1) plus one novel sensory neuron type with cell bodies in the proboscis labellum (diatom) (Figure 4 – figure supplement 2). Cell types in this group arborize in the anterior and superior region of the GNG, inferior to Group 1. Of the 18 members of this group, only one cell type, ‘feeding neuron’ (Fdg), has been previously described (Flood et al., 2013). Fdg neurons respond to food presentation in starved flies and activation of Fdg induces a feeding sequence. The split-GAL4 lines reported here greatly improve genetic access to Fdg, which was previously identified via stochastic activation of neurons labeled by a broadly expressed GAL4 line (Flood et al., 2013). Several other Group 2 neurons (indigo, tinctoria) are located near previously identified pumping motor neurons (Manzo et al., 2012; McKellar et al., 2020). This proximity, considered alongside the inclusion of Fdg and diatom, may suggest that other Group 2 neurons have roles in feeding sequence generation.

**Figure 4.**
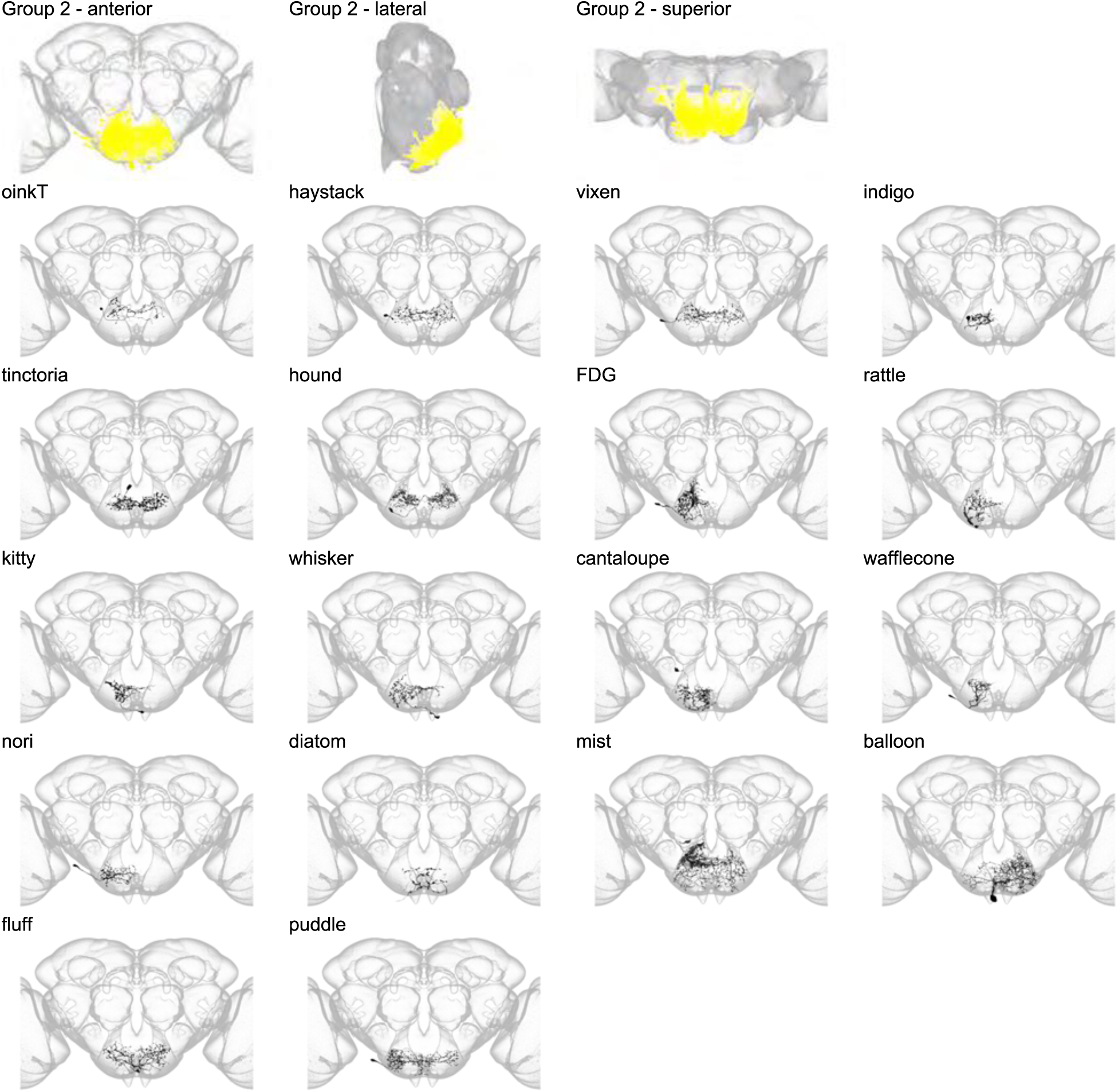
Morphology of individual neuron types in Group 2. The top row shows the morphology of all neuron types in Group 2 (yellow), with the morphology of individual group members shown below.

Group 3 contains 27 members and is composed of SEZ projection neurons and interneurons that overlap with the dendrites of these projection neurons (Figure 5 and Figure 5 – figure supplement 1). In the SEZ, Group 3 neurons arborize just anterior to the boundary between the anterior and posterior SEZ and in the superior region of the GNG, sometimes innervating the SAD or vest. Group 3 sits posterior to both Groups 1 and 2 in the SEZ. In contrast to Group 1 projection neurons, Group 2 projection neurons innervate diverse brain regions, including the superior lateral protocerebrum, superior clamp, and posteriorlateral protocerebrum, among others (Figure 9). One member of Group 3 has been previously reported, gustatory second-order neuron type 1 (G2N-1) (Miyazaki et al., 2015). The presence of G2N-1 in Group 3 and the proximity of this group to gustatory receptor neuron axons indicates that some Group 3 interneurons may participate in taste processing. Furthermore, Group 3 projection neurons innervate recently described regions of the SLP that respond to taste stimulation (Kim et al., 2017; Snell et al., 2020). Thus, Group 3 may in part be composed of taste responsive neurons.

**Figure 5.**
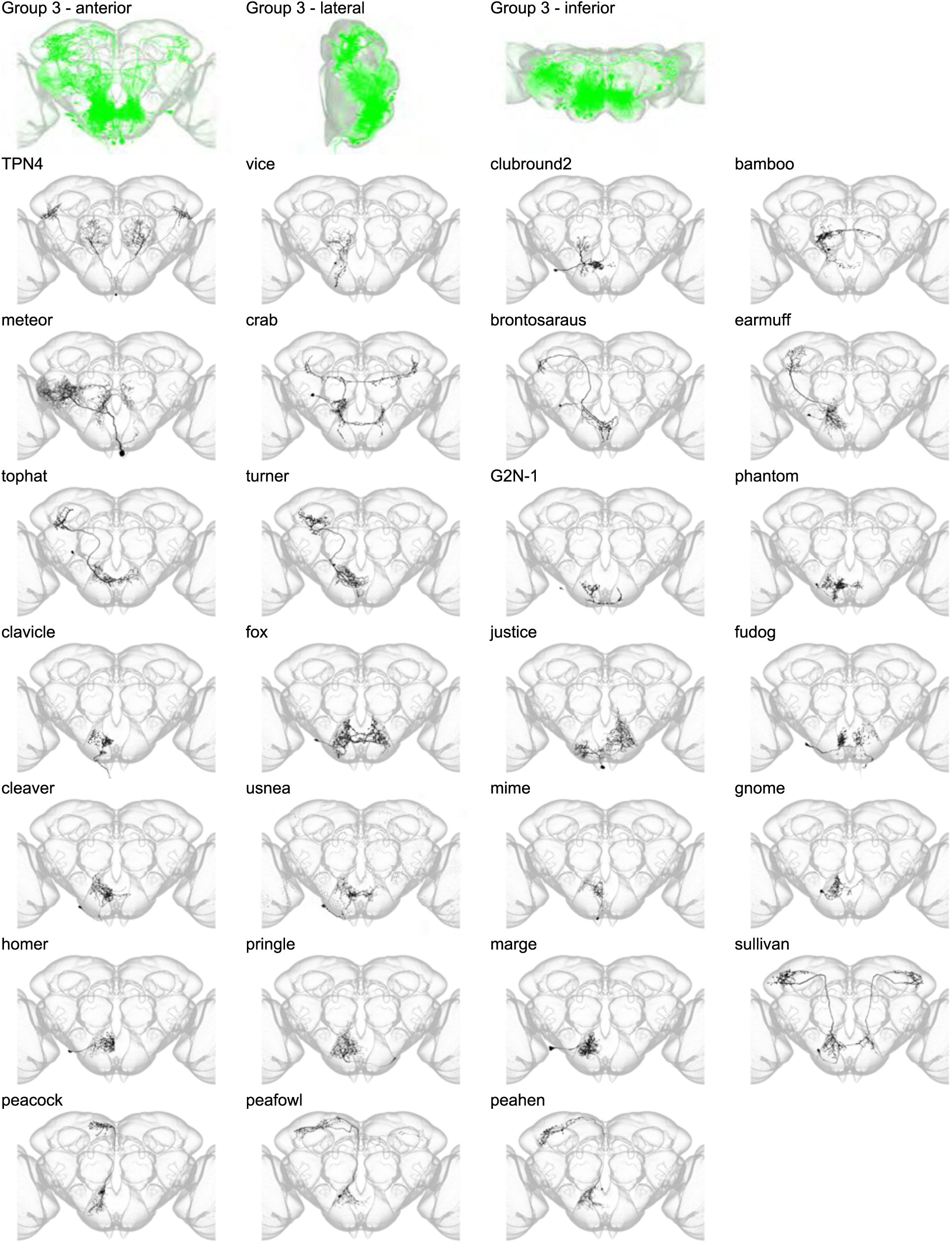
Morphology of individual neuron types in Group 3. The top row shows the morphology of all neuron types in Group 3 (green), with the morphology of individual group members shown below.

Group 4 contains interneurons that arborize in the inferior GNG and wrap the inferior surface of the GNG (Figure 6). Group 4 sits inferior to Groups 2 and 3. Notably, Group 4 includes five novel DNs that arborize in the GNG and descend to the leg neuropil (LegNp) in the prothoracic neuromere (Figure 6 – figure supplement 1) (Court et al., 2020). Of the 26 members of Group 4, only one, tyrosine hydroxylase ventral unpaired medial (TH-VUM) has been previously reported. TH-VUM is a dopaminergic neuron that influences the probability of proboscis extension (Marella et al., 2012). Notably, many interneurons in Group 4 are located near proboscis motor neurons that control rostrum protraction, haustellum extension, and labellar spreading (Kendroud et al., 2018; McKellar et al., 2020). This proximity, when considered with the inclusion of DNs that innervate the leg neuropil, may indicate that Group 4 members function in proboscis motor control and coordination of feeding with locomotion.

**Figure 6.**
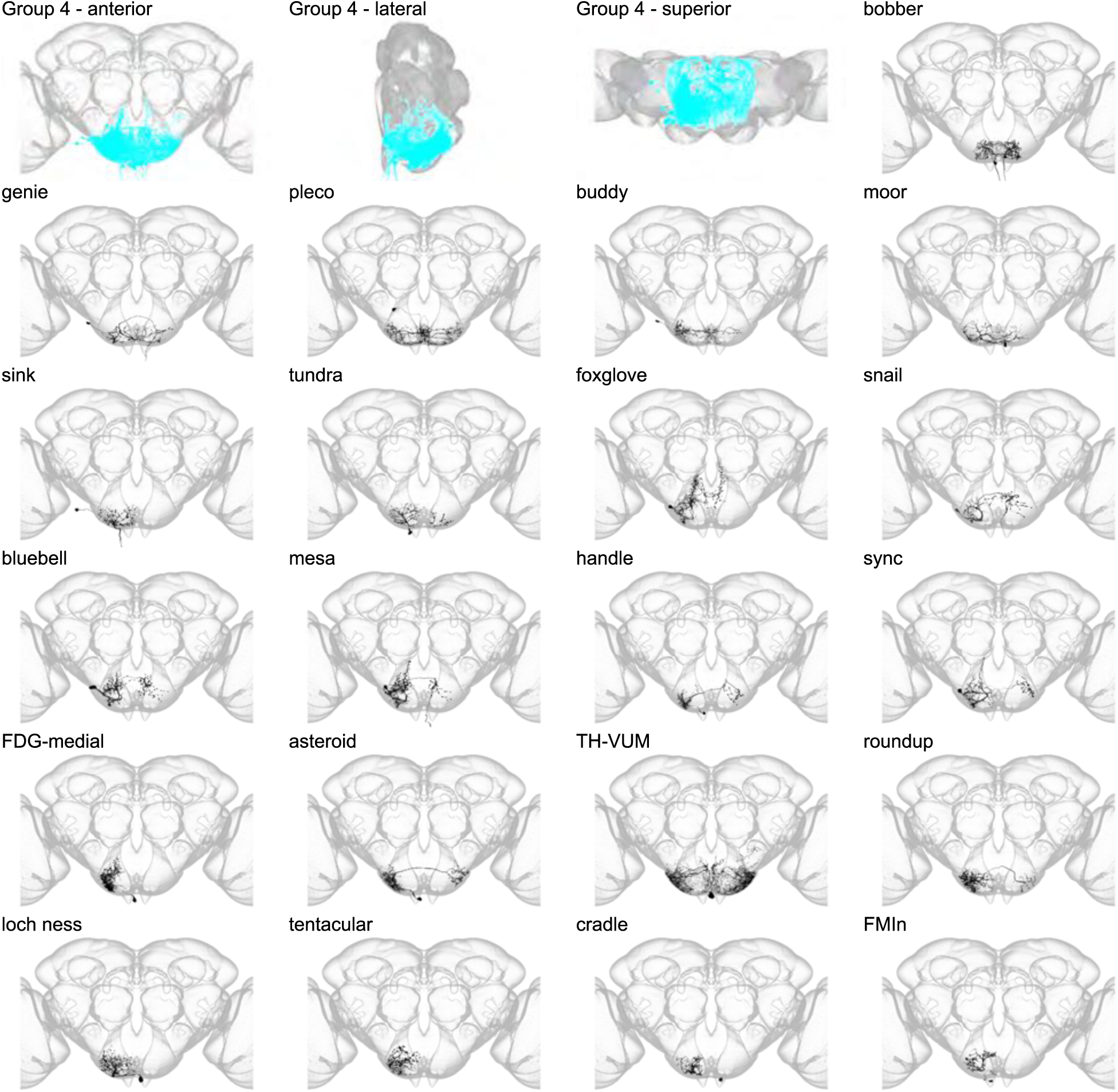
Morphology of individual neuron types in Group 4. The first three panels show the morphology of all neuron types in Group 4 (cyan), with the morphology of individual group members shown below.

Group 5 contains 27 interneurons, projection neurons, and DNs that arborize just posterior to the boundary between the anterior and posterior SEZ, flanked by Groups 3, 4, and 6 on the anterior, inferior, and posterior sides, respectively (Figure 7). Projection neurons in Group 5 arborize in the lobula, inferior and superior clamp, and inferior bridge, among other regions (Figure 9). The seven DNs in this group send their axons most frequently to the leg neuropils and to the abdominal ganglion (Abd) (Figure 7 – figure supplement 1). Three members of Group 5 have been previously reported. Two are previously described Fru+ neurons, aSG1 and aSG4 (Yu et al., 2010). The third neuronal type, aBN1, triggers antennal grooming when activated (Hampel et al., 2015). The proximity of neurons in Group 5 to previously described stopping neuron MAN (Bidaye et al., 2014), and the inclusion of an antennal grooming neuron, suggest that Group 5 neurons may participate in circuits that control grooming and stopping behaviors.

**Figure 7.**
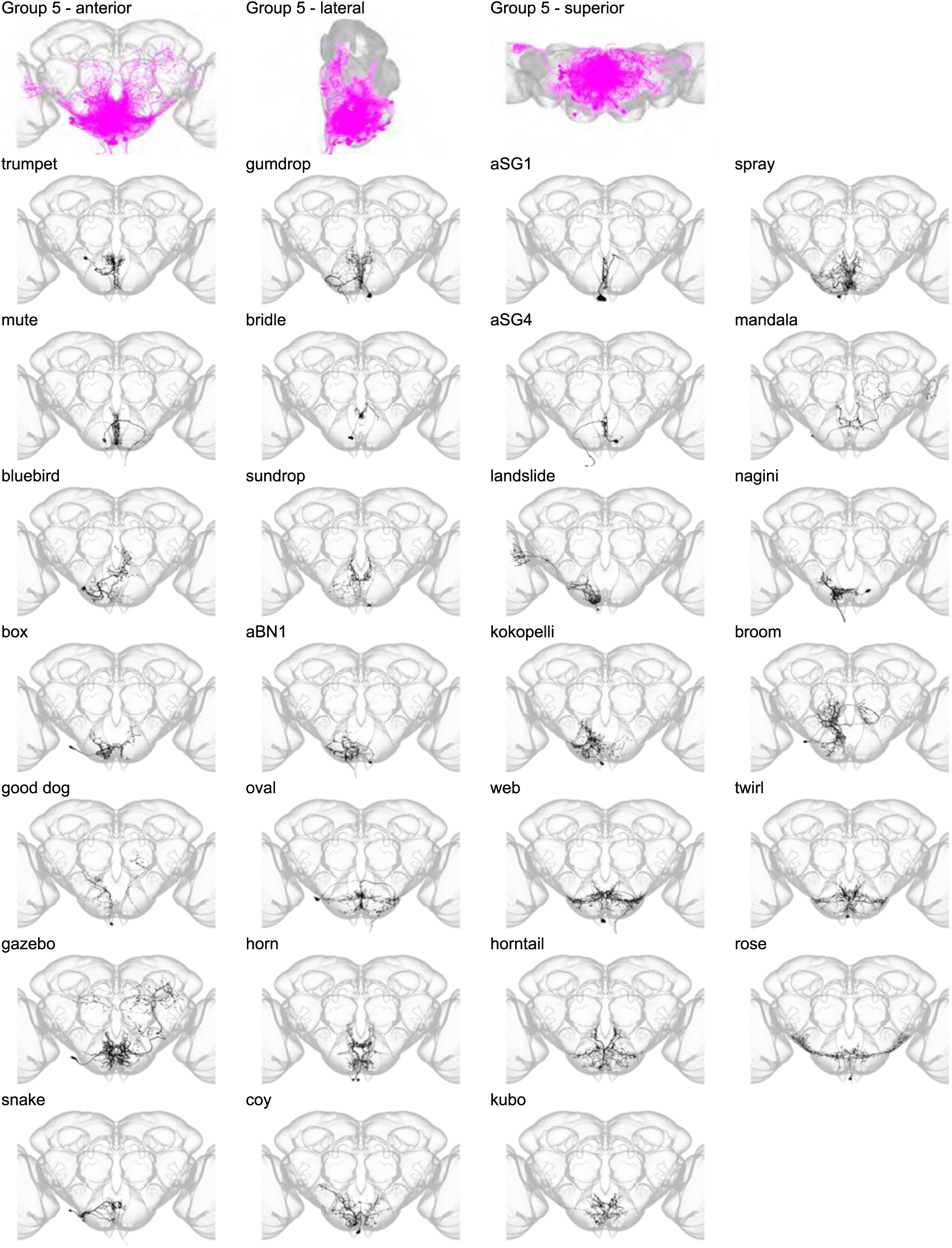
Morphology of individual neuron types in Group 5. The top row shows the morphology of all neuron types in Group 5 (fuchsia), with the morphology of individual group members shown below.

Group 6 contains neurons in the posterior of the brain, spanning the GNG, inferior posterior slope, and superior posterior slope (Figure 8) and is the most posterior group in the SEZ. This small group contains only nine members, and Group 6 cell types do not project to higher neuropils. One member, dubbed knees, is a DN that innervates neck neuropil and wing neuropil. No members of this morphological group have been previously reported. Several previous studies have implicated the SEZ and the posterior slope in flight behaviors, including wing and neck control (Namiki et al., 2018; Robie et al., 2017). We speculate that members of Group 6 may participate in circuits that control these behaviors.

**Figure 8.**
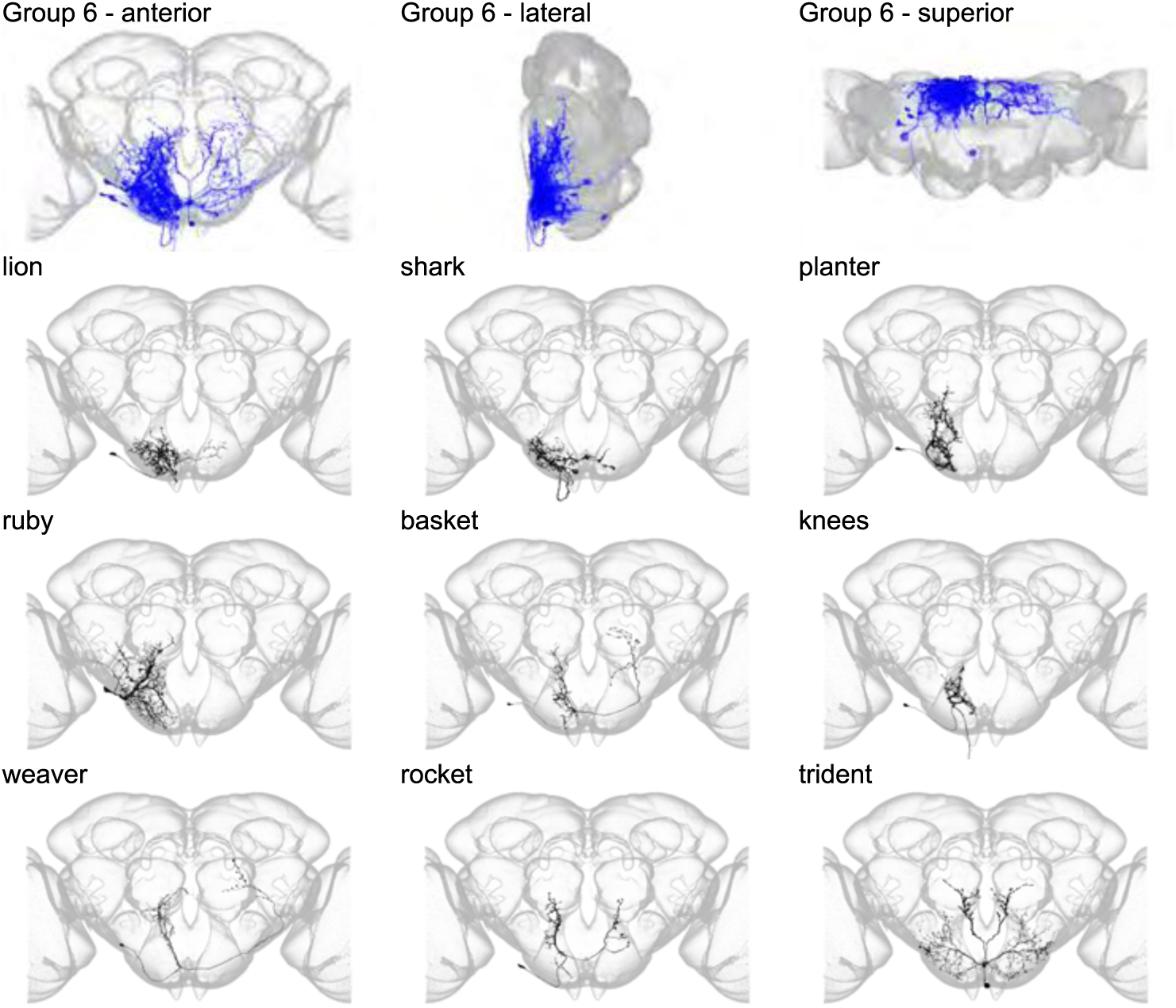
Morphology of individual neuron types in Group 6. The top row shows the morphology of all neuron types in Group 6 (royal blue), with the morphology of individual group members shown below.

Together, these six groups reveal layered and stacked structural domains in the SEZ. Whether neurons within the same morphological groups have similar functions remains unknown. However, this question may now be examined with the SEZ Split-GAL4 Collection, which provides reagents with which to evaluate how SEZ structure relates to function.

### SEZ interneurons tend to have mixed polarity

To shed light on the structure of information flow both within the SEZ and out of the SEZ to the higher brain and ventral nerve cord, we undertook polarity analysis of the 121 SEZ cell types that were segmented for NBLAST clustering analysis. These 121 cell types include 82 interneuron cell types, 26 projection neuron cell types, 12 DN cell types, and 1 sensory neuron cell type. We used both polarity staining with pre-synaptically localized HA-tagged Synaptotagmin and the smooth versus varicose appearance of neurites to score the presence of pre- and post-synaptic processes in each brain region in the central brain and ventral nerve cord (Court et al., 2020; Ito et al., 2014; Namiki et al., 2018). Upon examination of many cell types, we found that SEZ cell types frequently lack a defined axon and dendrite. Instead, inputs and outputs are mixed on the same processes. We designated these cell types as possessing “mixed” polarity. Other cell types have mostly mixed polarity but still retain a distinct arbor region where synaptic outputs are concentrated. We termed this category of cell types to have “biased” polarity. A third category is classically “polarized” with clearly separated processes dedicated to either synaptic input or synaptic output. To supplement our annotation of the presence of axons, dendrites, or both in each neuropil compartment and polarization strategy, we also indicated whether each cell type is an interneuron, projection neuron, DN, or sensory neuron (Figure 9, left).

**Figure 9.**
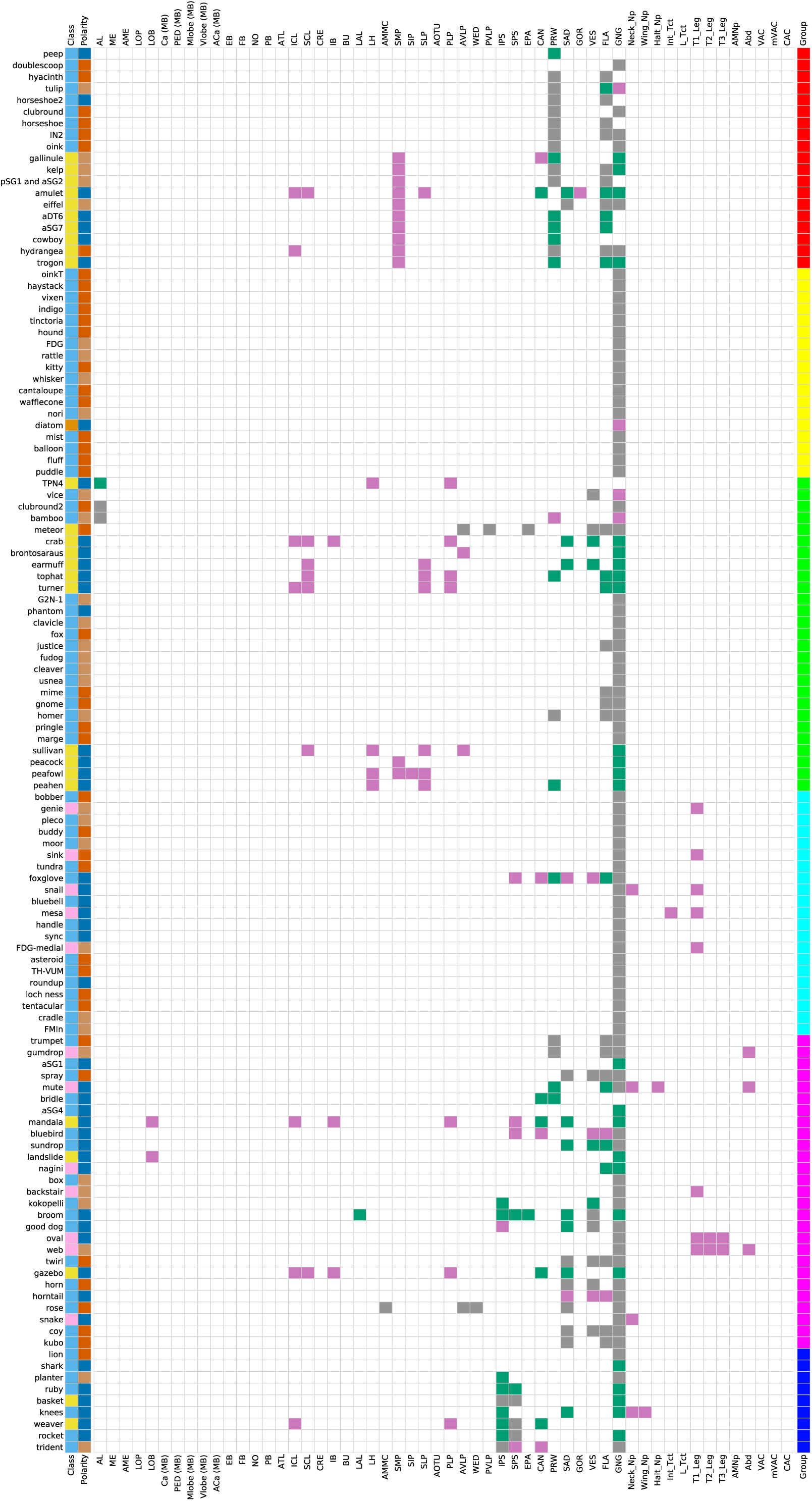
Innervation profile of SEZ neuron types. (Left-most column) Cell types are members of one of four cell type classes: interneuron (light blue), projection neuron (yellow), descending interneuron (light pink), or sensory neuron (orange). Interneurons are confined within the SEZ, while projection neurons project from the SEZ to higher neuropils in the central brain. Descending interneurons project their axons from the SEZ through the neck connective to the VNC. Sensory neurons project their axons from elsewhere in the body into the SEZ. Class is indicated for each neuron type by the filled pixel to the right of each neuron type name. (Second-to-leftmost column) Neuron types are polarized in a biased (light brown), mixed (red-orange), or polarized (dark blue) manner. The polarity strategy for each neuron type is indicated. (Center) The innervation profile for each neuron type is indicated by the filled pixels in its corresponding row. Brain region abbreviations follow the definitions and naming conventions of Ito et al. (2014) for the central brain and Court et al. (2020) for the VNC. The location of smooth processes (dendrites, green), varicose processes (axons, dark pink) or both smooth and varicose processes (axons and dendrites, grey) are indicated by defined neuropil region. VNC neuropil regions are grouped on the right of the figure. Innervation of the VNC was varicose in all cases. (Far right) Cell group as defined by NBLAST clustering is indicated for each cell type. Group 1, red; Group 2, yellow; Group 3, green; Group 4, cyan; Group 5, fuchsia; Group 6, royal blue.

Among the SEZ interneuron cell types we analyzed, 40/82 (49%) have mixed polarity, 22/82 (27%) have biased polarity, and 20/82 (24%) are classically polarized. Interneuron types are distributed throughout the six cell type supergroups with most groups containing interneurons of all polarity classes. However, all Group 2 interneurons (making up 17/18 cell types in Group 2) have either biased or mixed polarity. This suggests that the interneurons of Group 2 may participate in reciprocally connected circuits. Among interneuron cell types that are clearly polarized, there were some cases in which no axon was evident in the brain (including peep, shark, bridle, aSG1, and aSG4). In these cases, the presence of severed processes suggests that these cell types may not be interneurons and may instead send projections out of the central nervous system.

Most SEZ projection neurons analyzed in this study are clearly polarized (20/26, 77%). However, a few have biased (4/26, 15%) or mixed polarity (2/26, 8%). Polarized cell types belong to several cell type Groups (3, 5, 6) and project to numerous brain regions including the lobula and the superior, inferior, and ventrolateral neuropils. In contrast, all projection neuron cell types with biased polarity have axons in the SMP and belong to Group 1. The high proportion of clearly polarized SEZ projection neurons suggests that they commonly carry information uni-directionally from the SEZ to other brain regions. We did not identify any projection neurons that link the SEZ directly to the central complex or the mushroom body. In addition to projection neurons, the SEZ Split-GAL4 Collection includes one novel sensory neuron class (diatom) that is polarized, with dendrites in the proboscis labellum and SEZ axonal projections (Figure 4 and Figure 4 – figure supplement 1).

Analyzing the polarity of the novel SEZ DNs described here, we find that 5/12 (42%) have biased polarity, 1/12 (8%) has mixed polarity, and 6/12 (50%) are clearly polarized. These DNs have outputs in the neck, haltere, and leg neuropils (in all three neuromeres), and in the abdominal ganglion, consistent with the observation that GNG DNs are frequently connected with the leg neuropils (Namiki et al., 2018).

Thus, polarity analysis reveals that most types of SEZ interneurons have either mixed or biased polarity. In contrast, SEZ projection neurons are frequently clearly polarized while the SEZ DNs reported here are mostly mixed or polarized. These distinct polarization strategies may reflect the functional roles that interneurons, projection neurons, and DNs play in SEZ circuits. Interneurons in the SEZ may participate in reciprocally connected local networks while projection neurons and DNs may primarily relay information from the SEZ to other brain regions.

## Discussion

Here, we describe the SEZ Split-GAL4 Collection, a library of 283 split-GAL4 lines covering 138 SEZ cell types, which affords unprecedented genetic access to SEZ neurons for behavioral and functional study. The SEZ Split-GAL4 Collection provides insight into the anatomical organization of the SEZ and a resource with which to investigate how diverse sensory inputs, including gustatory, pheromone, and mechanosensory inputs, are processed by local SEZ circuitry and ascending paths to control specific behaviors.

Most of the SEZ Split-GAL4 lines are specific, with 150/283 lines classified as ideal or excellent. These lines will be useful to manipulate individual SEZ cell types for behavioral, functional, and imaging experiments. The remaining, less specific, lines (those belonging to the good or poor categories) will still be useful for imaging and as starting points for creating more specific reagents. Good and poor lines may be used to generate CDM masks to search for new hemidrivers to make further split-GAL4 lines. Alternatively, their expression patterns may be refined using Killer Zipper or three-way intersections with LexA or QF lines (Dolan et al., 2017; Shirangi et al., 2016). All lines in the SEZ Split-GAL4 Collection may be used to generate further tools including complementary split-LexA and split-QF reagents (Riabinina et al., 2019; Ting et al., 2011). Split-LexA and split-QF lines may be used in concert with the split-GAL4 lines reported here to simultaneously manipulate two independent neuronal populations for advanced intersectional experiments, including behavioral epistasis.

By combining insights from a single-cell transcriptome atlas with direct cell counts of SEZ neuromeres, we estimate that the SEZ Split-GAL4 Collection labels 30% of the ∼1700 neurons in the SEZ. Our estimate of SEZ neuron number is based on analysis of the four genetically defined SEZ neuromeres, the tritocerebral, the mandibular, the maxillary, and the labial, and does not include potential deutocerebral contributions (Boyan et al., 2003). We emphasize that our estimate of SEZ neuron number is likely to be superseded by counts of SEZ neuronal cell bodies in EM volumes. Regardless, the SEZ Split-GAL4 Collection targets 510 neuronal cell bodies, which represents a substantial improvement in our ability to precisely target SEZ cell types for functional and morphological analysis. Furthermore, we did not ascertain the neuromere or neuroblast of origin of the SEZ cell types in the SEZ Split-GAL4 Collection. Recent work has established reliable anatomical criteria that define the boundaries between the four SEZ neuromeres and has mapped all secondary lineages of the SEZ (Hartenstein et al., 2018). Future efforts should focus on bridging previously identified fascicle, neuropil, and sensory domains into a common template or coordinate space to determine the neuromere and neuroblast origin of SEZ cell types.

Discovering and genetically targeting SEZ cell types required the use of registered light-level imagery and computer-assisted searching. We used four distinct strategies to identify 129 novel and 9 previously reported SEZ cell types in registered light-level imagery. Critically, each of these strategies allowed us to use CDM mask searching to identify additional hemidrivers with which to target each cell type of interest. CDM mask searching enabled combing of large datasets and greatly increased the ease and speed of split-GAL4 generation over previous methods (Otsuna et al., 2018). The same strategies can be leveraged to gain genetic access to yet-undiscovered SEZ cell types. The recent electron microscopy (EM) volumes of the *D. melanogaster* brain provide an avenue for identifying SEZ cell types that are not covered by the SEZ Split-GAL4 Collection. Notably, this approach awaits comprehensive reconstruction of the SEZ, a region that is not included in the recently published dense reconstruction of the “hemibrain” volume (Scheffer et al., 2020). Another EM volume, “FAFB”, provides imagery of an entire adult female fly brain at synaptic resolution, and includes the SEZ (Zheng et al., 2018). Improvements in automated reconstruction of EM volumes coupled with large-scale human annotation should soon provide exhaustive reconstruction of the SEZ from which to identify additional SEZ cell types (Dorkenwald et al., 2020). Furthermore, available bridging registrations between EM volumes and light level imagery should facilitate the identification of hemidrivers to target SEZ cell types discovered from EM reconstructions (Bates et al., 2020). Even without identifying additional SEZ cell types, the split-GAL4 reagents described will allow behavioral and functional evaluation of circuit hypotheses derived from EM imagery.

Our analyses of the SEZ Split-GAL4 Collection provide insight into the cellular architecture of the SEZ. To computationally probe the organization of the SEZ, we morphologically clustered 121 SEZ cell types using NBLAST (Costa et al., 2016). This approach reveals six cellular domains in the SEZ which are organized in a largely layered fashion from anterior to posterior. This layered structure is also hinted at by the recent description of SEZ neuropil domains throughout development from the larva to the adult (Kendroud et al., 2018). Based on anatomical position and the known function of a few SEZ neurons, we hypothesize that different morphological clusters may participate in different behavioral functions.

For example, Group 1 neurons arborize in the tritocerebrum and SMP and may participate in regulation of nutritive state, Groups 2-4 may participate in feeding, taste processing and proboscis motor control, and central Group 5 and more posterior Group 6 may be involved in grooming and flight motor control. While currently speculative, these hypotheses can now be rigorously tested by querying the behavioral and physiological function of SEZ neurons with exquisite precision using the split-GAL4 reagents reported here. The combination of specific access to SEZ cell types and detailed connectivity maps from EM studies now provides a path to elucidate the organizational principles of the SEZ.

Our studies also shed light on information flow both within the SEZ and out of the SEZ to the higher brain. Polarity analysis of 121 SEZ cell types revealed that SEZ interneurons tend to have mixed or biased polarity while SEZ projection neurons tend to be classically polarized. This distinction suggests that the interneurons that compose local SEZ circuits are highly reciprocally interconnected, with information flowing in networks rather than unidirectional streams. Projection neurons, in contrast, may serve chiefly to pass information from highly interconnected SEZ circuits to other brain regions in a unidirectional manner. Notably, we identified many SEZ projection neurons that innervate the SMP—a region known to contain neurosecretory cell types. This may betray a role for acute taste detection or feeding circuit activation in the regulation of hormone secretion. In addition, the frequent innervation of the superior lateral protocerebrum and lateral horn by SEZ projection neurons may hint at the site of olfactory-gustatory synthesis. In contrast, we did not identify projection neurons that link the SEZ directly to the central complex or mushroom body. If dense reconstruction of EM volumes corroborates the lack of direct connectivity between the SEZ and these regions, information must be conveyed through indirect pathways. As an example, taste information influences local search behaviors during foraging, a task that is expected to involve the central complex (Haberkern et al., 2019). Indirect relay of taste information to the central complex to inform foraging behavior would be consistent with previous anatomical studies suggesting that the central complex receives diverse indirect sensory inputs (Pfeiffer and Homberg, 2014). Furthermore, the mushroom body is known to respond to taste, raising the possibility that taste information from gustatory sensory neuron axons in the SEZ must be relayed through yet another brain region before reaching mushroom body cell types (Harris et al., 2015). Thus, our analysis of SEZ neuron polarity indicates local SEZ processing and demonstrates direct pathways to a subset of higher brain regions.

Overall, the SEZ Split-GAL4 Collection represents a valuable resource that will facilitate the study of the SEZ. Our analysis of the collection reveals the cellular anatomy and polarity of individual SEZ neurons and their organization into six discrete domains in the SEZ. Coupled with emerging insights from reconstruction of EM volumes, the SEZ Split-GAL4 Collection will allow the use of genetic dissection to test circuit-level hypotheses about sensory processing and motor control in the SEZ.

## Materials and Methods

### Key Resources Table

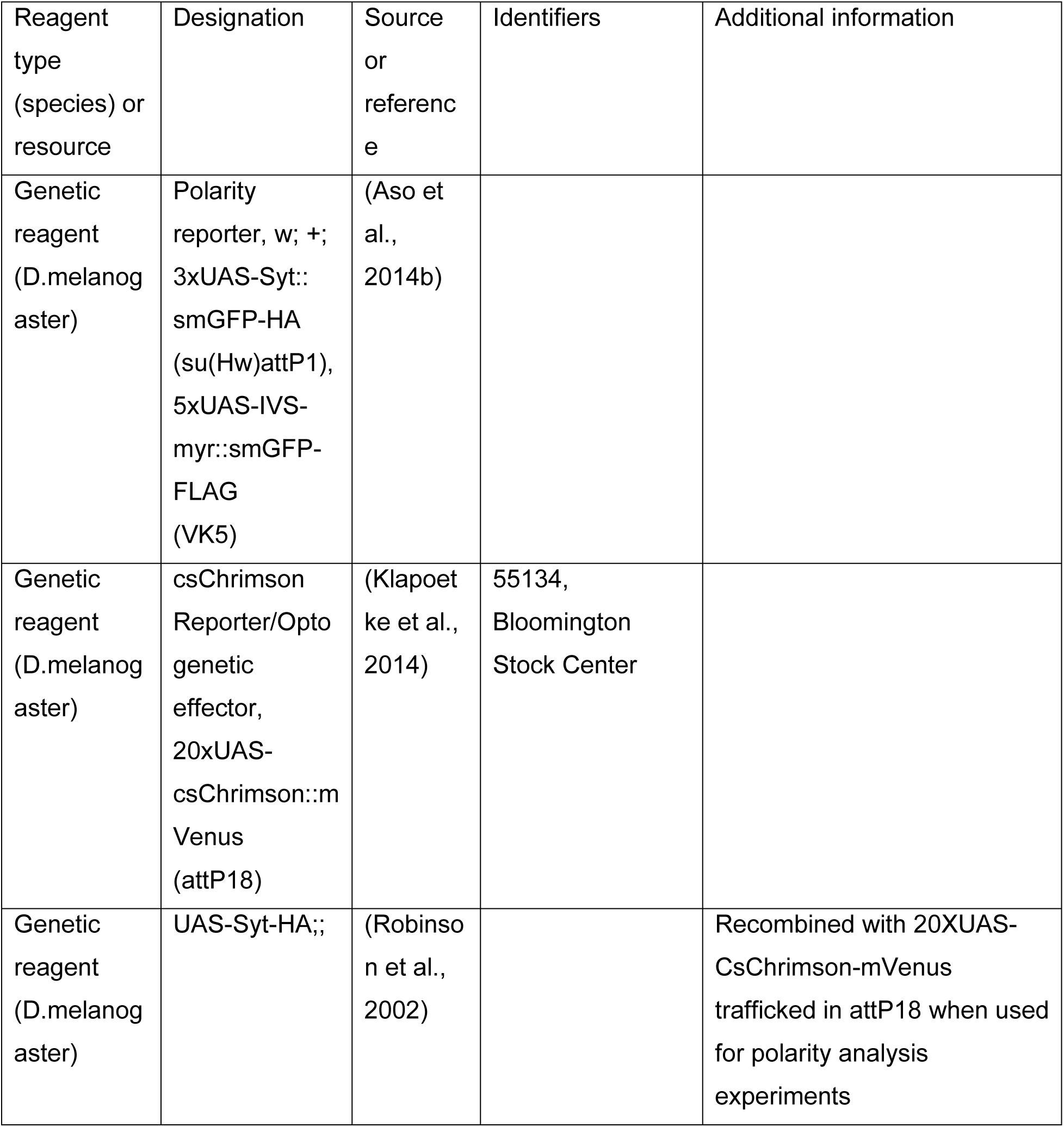
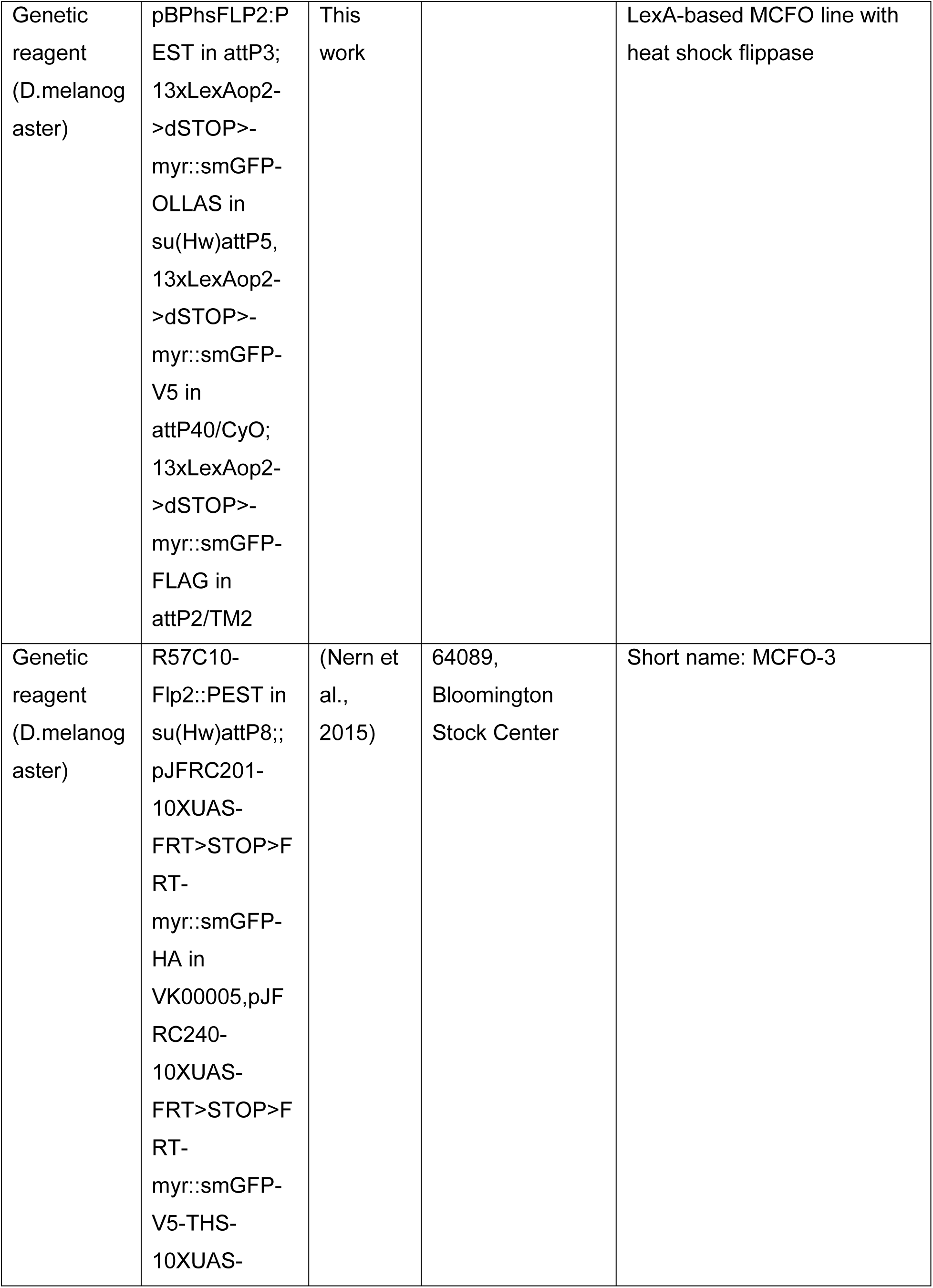
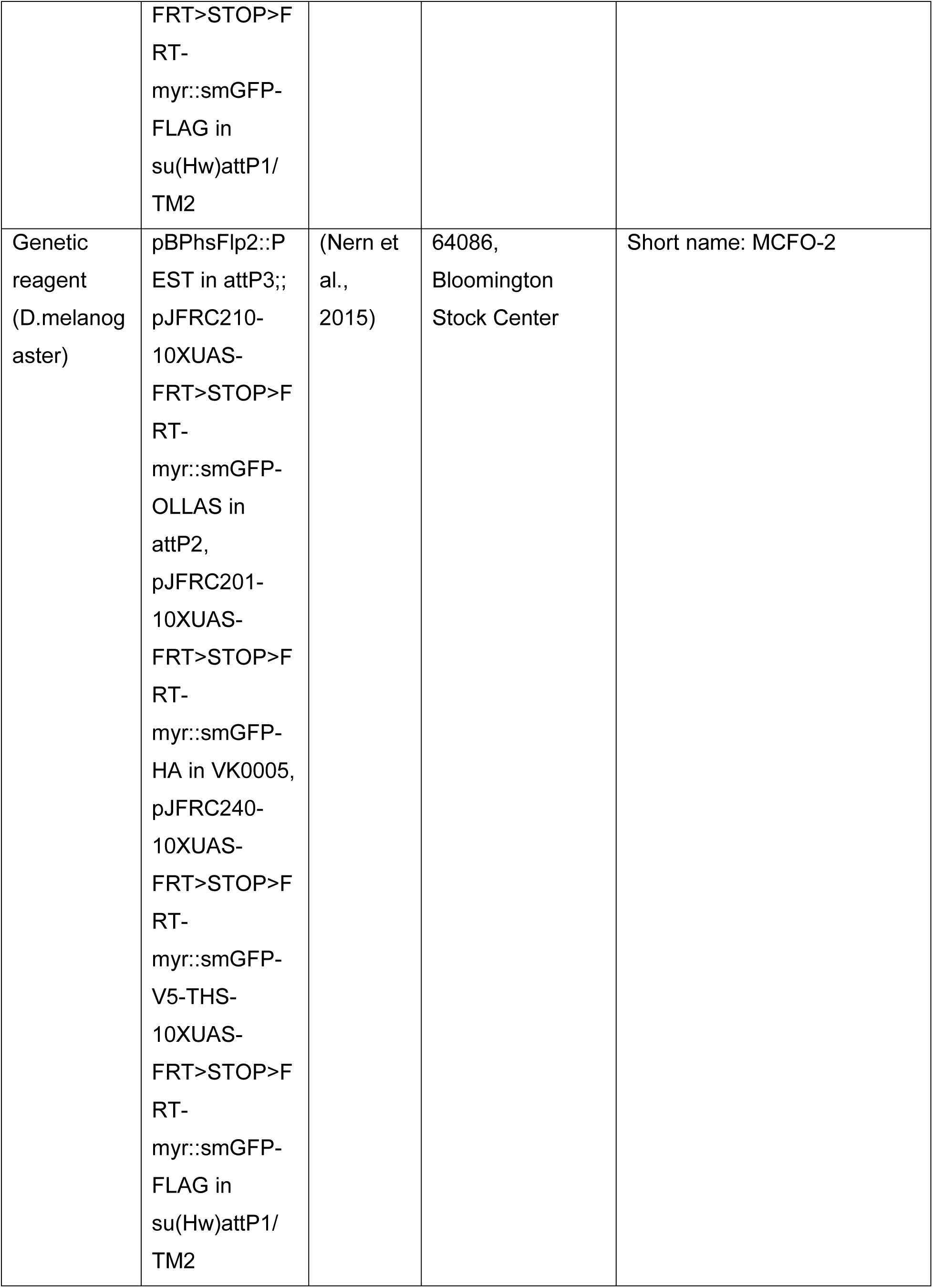
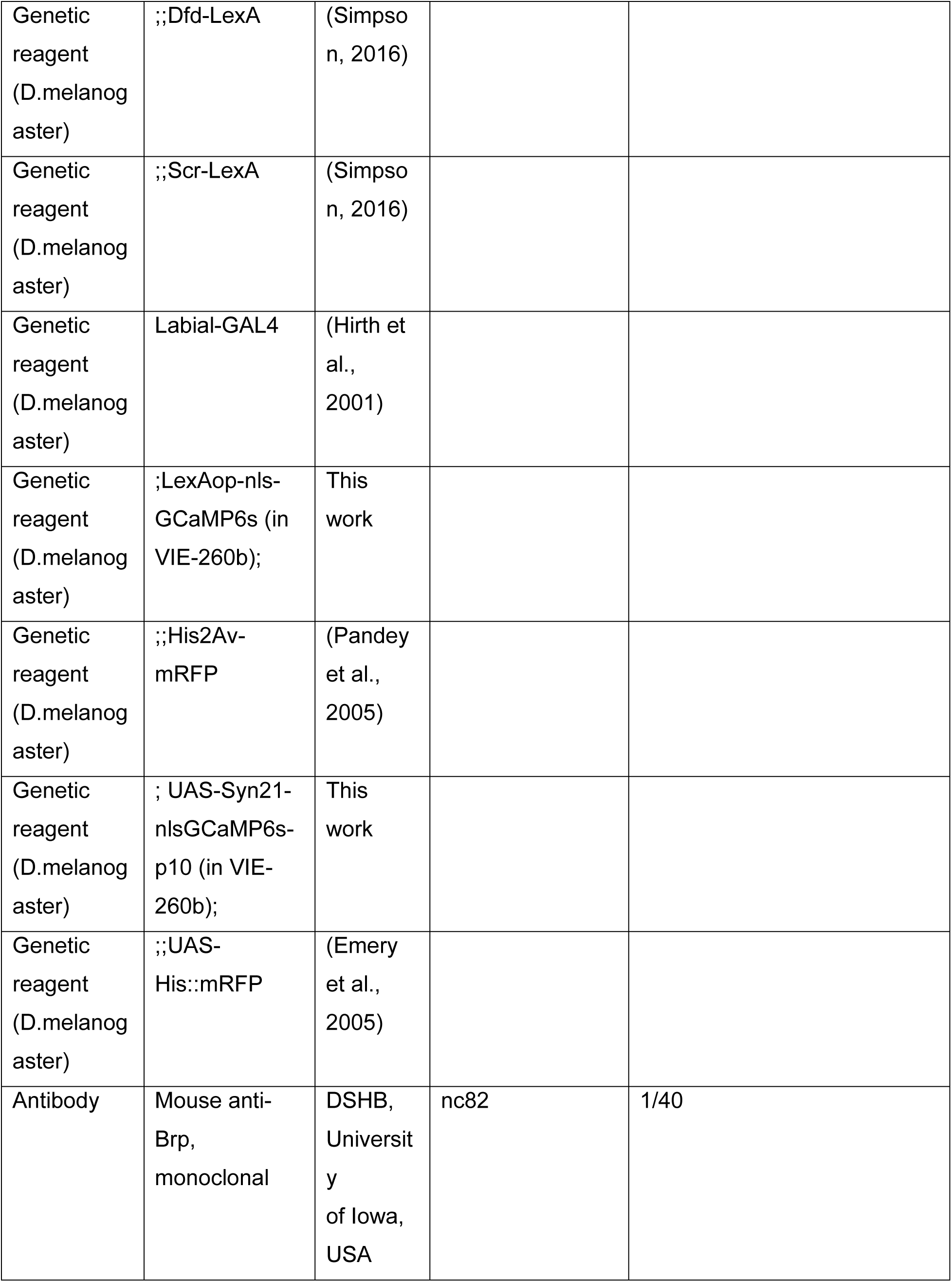
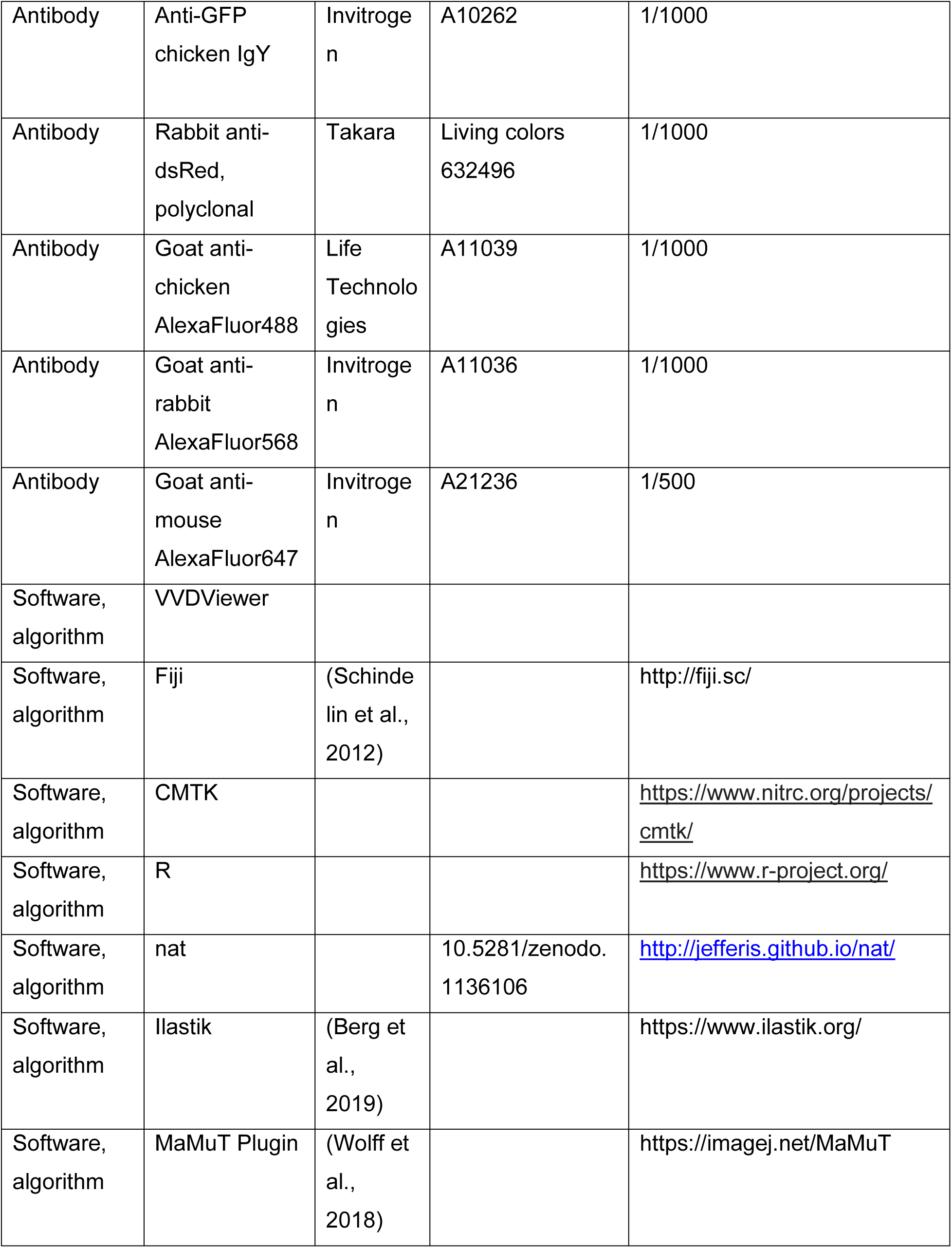
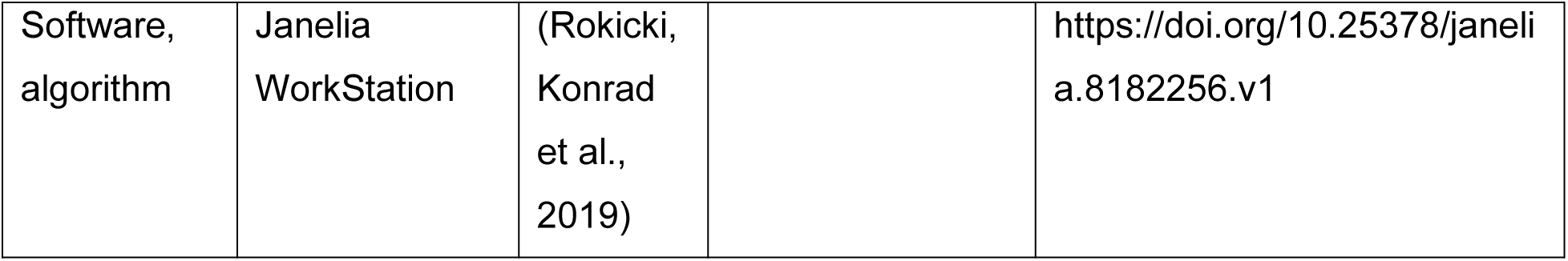

### Drosophila Husbandry

All experiments and screening were carried out with adult *Drosophila melanogaster* females raised at 25°C on standard *Drosophila* food. Adult females were mated and dissected within one week of eclosion. Construction of stable split-GAL4 lines was performed as previously described (Dionne et al., 2018).

### Counting SEZ Neurons

Either Dfd-LexA or Scr-LexA was crossed to a reporter line with LexAop-nls-GCaMP6s (this work) and His2Av-mRFP (Pandey et al., 2005). Labial-GAL4 was crossed to a reporter line with UAS-nls-GCaMP6s (this work) and UAS-His2Av-mRFP (Emery et al., 2005). Brains dissected as described (https://www.janelia.org/project-team/flylight/protocols, “Dissection and Fixation 1.2% PFA”).

The following primary antibodies were used:

- 1/40 Mouse α-Brp (nc82) (DSHB, University of Iowa, USA)
- 1/1000 Chicken α-GFP (Invitrogen A10262)
- 1/1000 Rabbit α-dsRed (Takara, Living Colors 632496)

The following secondary antibodies were used:

- 1/500 α-mouse AF647 (Invitrogen, A21236)
- 1/1000 α-chicken AF488 (Life Technologies, A11039)
- 1/1000 α-rabbit AF568 (Invitrogen, A21236)

Immuhistochemistry was carried out as described (https://www.janelia.org/project-team/flylight/protocols, “IHC - Anti-GFP”) substituting the above antibodies and eschewing the pre-embedding fixation steps. Ethanol dehydration and DPX mounting was carried out as described (https://www.janelia.org/project-team/flylight/protocols, “DPX Mounting”). Images were acquired with a Zeiss LSM 880 NLO AxioExaminer at the Berkeley Molecular Imaging Center. A Plan-Apochromat 63x/1.4 Oil DIC M27 objective was used at zoom 1.0. Acquired images had a voxel size of 0.132 μm x 0.132 μm x 0.500 μm. Expression in the SEZ was imaged in a tiled fashion and then stitched in Fiji using the “Grid/Collection stitching” plugin with “Unknown Positions” and “Linear Blending” (Preibisch et al., 2009).

SEZ cell number was quantified with Ilastik using the “Pixel Classification” and “Object Classification” workflows (Berg et al., 2019). The pixel classifier was trained to segment only cell bodies expressing both LexAop-nls-GCaMP6s and His2Av-mRFP, which improved pixel and object classification accuracy when compared to using LexAop-nls-GCaMP6s without His2Av-mRFP (data not shown). Then, to verify counts derived from automated Ilastik quantification, manual ground truth counts of example image regions (four subregions each for Dfd-LexA and Scr-LexA) were compared to counts of the same regions derived from Ilastik. Ground truth counts were carried out in three dimensions with the MaMuT plugin in Fiji (Wolff et al., 2018). Error was calculated at 0.5% for Dfd-LexA images and -1.4% for Scr-LexA images.

### Split-GAL4 intersections

Novel SEZ cell types were identified using the following strategies:

1. Visual search through several large, publicly available GAL4 collections designed to tile the nervous system (Jenett et al., 2012; Pfeiffer et al., 2008; Tirian and Dickson, 2017).
2. LexA-based MultiColor FlpOut (MCFO) of Scr-LexA and Dfd-LexA. In total, 232 Scr-LexA samples and 320 Dfd-LexA samples were examined.
3. MCFO (Nern et al., 2015) screening of subsets of the Janelia Research Campus and Vienna Tile GAL4 collections that have dense SEZ expression in which individual cell morphologies were difficult to parse (Meissner et al., 2020). 66,080 CDM images from MCFO of 2182 unique lines were examined.
4. Re-registration of open-access images of individual SEZ cell types from MARCM screens of broad GAL4 drivers, available on FlyCircuit (Chiang et al., 2011). 22,598 female samples re-registered to the “JFRC 2010” template (Jenett et al., 2012) were analyzed.

Following identification of cell types, we created representative CDM masks and used CDM mask searching (Otsuna et al., 2018) to find additional enhancers whose expression patterns seemed to include the desired cell type. We annotated all drivers that putatively drove expression in each of the identified cell types. We searched the following CDM images: 27,534 CDM images covering 6575 Janelia Research Campus GAL4 lines; 18,047 CDM images covering 8031 GAL4 Vienna Tile lines; and 66,080 CDM images from MCFO of 2182 unique Janelia Research Campus and Vienna Tile lines. In total we used 86,861 CDM images for CDM mask searching. We then assessed the availability of hemidrivers for each of the enhancers (ADs and DBDs). The split-GAL4 hemidrivers used in this study were previously generated at Janelia Research Campus (Dionne et al., 2018; Tirian and Dickson, 2017). Then, the expression patterns for all possible AD-DBD combinations for a given cell type were screened. Screening was carried out in adult female flies as previously described (Dionne et al., 2018). A single female central nervous system was screened per combination. With few exceptions, screening was carried out by FlyLight using the FLylight split-screen protocol: (https://www.janelia.org/project-team/flylight/protocols, “IHC - Adult Split Screen”). Following dissection, staining, and mounting, split-GAL4 combinations were screened by eye using epifluorescence on an a LSM710 confocal microscope (Zeiss) with a Plan-Apochromat 20x/0.8 M27 objective. Imagery was viewed and organized using the Janelia Workstation (Rokicki, Konrad et al., 2019). Useful combinations with limited SEZ expression were selected for initial confocal imaging using a 20x objective. Following imaging, useful combinations were further sorted and annotated in a custom database. The resulting database of SEZ split-GAL4 lines contains the following:

- target cell type
- AD and DBD
- Unique JRC_SS identifier
- line quality (Ideal>Excellent>Good>Poor)
- a text description of any off-target expression
- types of imagery collected, including polarity and MCFO data

After stabilization (Dionne et al., 2018), select split-GAL4 lines were further characterized. We selected at least one split-GAL4 line per cell type for detailed documentation, including polarity staining (to assess expression pattern in multiple central nervous systems and to determine the location of synaptic outputs), MCFO characterization, and 63x imaging. Polarity staining was carried out by crossing stabilized split-GAL4 lines to either w; +; 3xUAS-Syt::smGFP-HA(su(Hw)attP1), 5xUAS-IVS-myr::smGFP-FLAG (VK5) or UAS-Syt-HA, 20XUAS-CsChrimson-mVenus (attP18);;. When crossed to w; +; 3xUAS-Syt::smGFP-HA(su(Hw)attP1), 5xUAS-IVS-myr::smGFP-FLAG (VK5) dissection and staining were carried out by FlyLight according to the FlyLight “IHC - Polarity Sequential” protocol: (https://www.janelia.org/project-team/flylight/protocols). When crossed to 20XUAS-CsChrimson-mVenus (attP18);; dissection and staining were carried out by FlyLight according to the FlyLight “IHC - Polarity Sequential Case 5” protocol: (https://www.janelia.org/project-team/flylight/protocols). MCFO characterization of stable split-GAL4 lines was accomplished by crossing stable lines to MCFO-2 or MCFO-3 (See “Key Resources Table” for full genotypes). If crossed to MCFO-2, adult flies were heat shocked at 37° C for either 30 or 60 minutes one day after eclosion. Dissection and staining of MCFO samples were carried out by FlyLight according to the FlyLight MCFO staining protocol: (https://www.janelia.org/project-team/flylight/protocols, “IHC - MCFO”). Samples stained for polarity and MCFO analysis were first imaged on an a LSM710 confocal microscope (Zeiss) with a Plan-Apochromat 20x/0.8 M27 objective. Then, sample images were viewed using the Janelia Workstation (Rokicki, Konrad et al., 2019) and several samples per line were chosen for higher resolution imaging. Higher resolution imaging of select samples was carried out on a LSM710 confocal microscope (Zeiss) with a Plan-Apochromat 63x/1.40 oil immersion objective. If multiple tiles were required to cover the region of interest, tiles were stitched together (Yu and Peng, 2011).

### Morphological clustering with NBLAST

63x MCFO images were registered to the full-size JRC 2018 unisex template (Bogovic et al., 2020) using CMTK (https://www.nitrc.org/projects/cmtk). A single example of each cell type targeted by the collection was selected for segmentation in VVD Viewer (https://github.com/takashi310/VVD_Viewer). The following 17 cell types covered by the SEZ split-GAL4 Collection were excluded because suitable MCFO images were not available: bay, bower, braces, bubbA, bump, clownfish, handup, linea, mothership, oinkU, pampa, portal, seagull, slink, spirit, stand, and willow. The expression pattern of the best split-GAL4 line for each excluded cell type is show (Figure 2 – figure supplement 1). The remaining 121 cell types covered by the collection were included in NBLAST analysis. Registration quality was assessed by viewing the overlap between the template brain and the registered nc82 reference channel to ensure that selected images were well registered. Further, selected images were only used if the morphology of the cell type of interest was clearly visible and not intermingled with other cells or neuronal processes that might lead to false merges or truncations due to neighboring cell types. Images were manually segmented in VVD Viewer to remove non-specific background and other, clearly distinct cells. Following segmentation, images were thresholded using the “Huang” method (Huang and Wang, 1995), flipped to the right hemisphere of the brain, and scaled to a final voxel size of (x) 0.3766 x (y) 0.3766 x (z) 0.3794. Scaled images were then skeletonized with the “Skeletonize 2d/3d” Fiji plugin (Lee et al., 1994). Skeletonized, scaled images were hierarchically clustered using NBLAST and Ward’s method (Costa et al., 2016). This was carried out with the natverse toolkit in R (Bates et al., 2020). Group number was chosen by assessing Ward’s joining cost and the differential of Ward’s joining cost after Braun et al. (2010). Images of the resulting morphological clusters were further visualized in R, again using natverse (Figure 2). Catalog figures were assembled using full-sized segmented imagery in VVD Viewer (Figures 3-8).

### Polarity Analysis

Full-size registered, segmented example neuron images (prior to scaling or skeletonizing) created as described above were compared against established neuropil regions (Court et al., 2020; Ito et al., 2014) in VVDViewer. The presence of smooth versus varicose processes were scored after Namiki et al. (2018). Images from polarity staining were referenced where available.

## Supporting information

SEZ Split-Gal4 Collection

## Funding

Howard Hughes Medical Institute (Janelia Research Campus)

- Gabriella R Sterne
- Barry J Dickson
- Hideo Otsuna

National Institute of Diabetes and Digestive and Kidney Diseases (F32DK117671)

- Gabriella R. Sterne

National Institute of General Medical Sciences (R01NS110060)

- Kristin Scott
- Gabriella R. Sterne

The funders had no role in study design, data collection and interpretation, or the decision to submit the work for publication.

## Acknowledgements

We would like to thank the Janelia Fly Facility (Todd Laverty, Amanda Cavallaro, Scarlett Harrison, Karen Hibbard, Jui-Chun Kao, and Guillermo Gonzalez among others) for help with fly husbandry and fly line generation. The FlyLight Project Team (https://www.janelia.org/project-team/flylight, Geoffrey Meissner, Kelley Lee, Zachary Dorman, Oz Malkesman, and others) performed brain dissections, immunohistochemistry, and confocal imaging for split-GAL4 screening and characterization after stabilization. We would like to acknowledge Geoffrey Meissner for testing and providing the recombined UAS-Syt-HA, 20XUAS-CsChrimson-mVenus trafficked in attP18 stock for split-GAL4 polarity analysis. Confocal imaging for the SEZ cell counting experiments were conducted at the CRL Molecular Imaging Center, supported the Helen Wills Neuroscience Institute and NSF DBI-1041078. We would like to thank Holly Aaron and Feather Ives for their microscopy training and assistance. Leo Guignard generously provided feedback and guidance without which automated cell counting with Ilastik would not have been possible. Ryo Minegishi shared code and tips for implementing NBLAST on new datasets. We would also like to thank Masayoshi Ito for sharing split-GAL4 combinations that labeled SEZ neurons.

---

**Figure 1—figure supplement 1.**
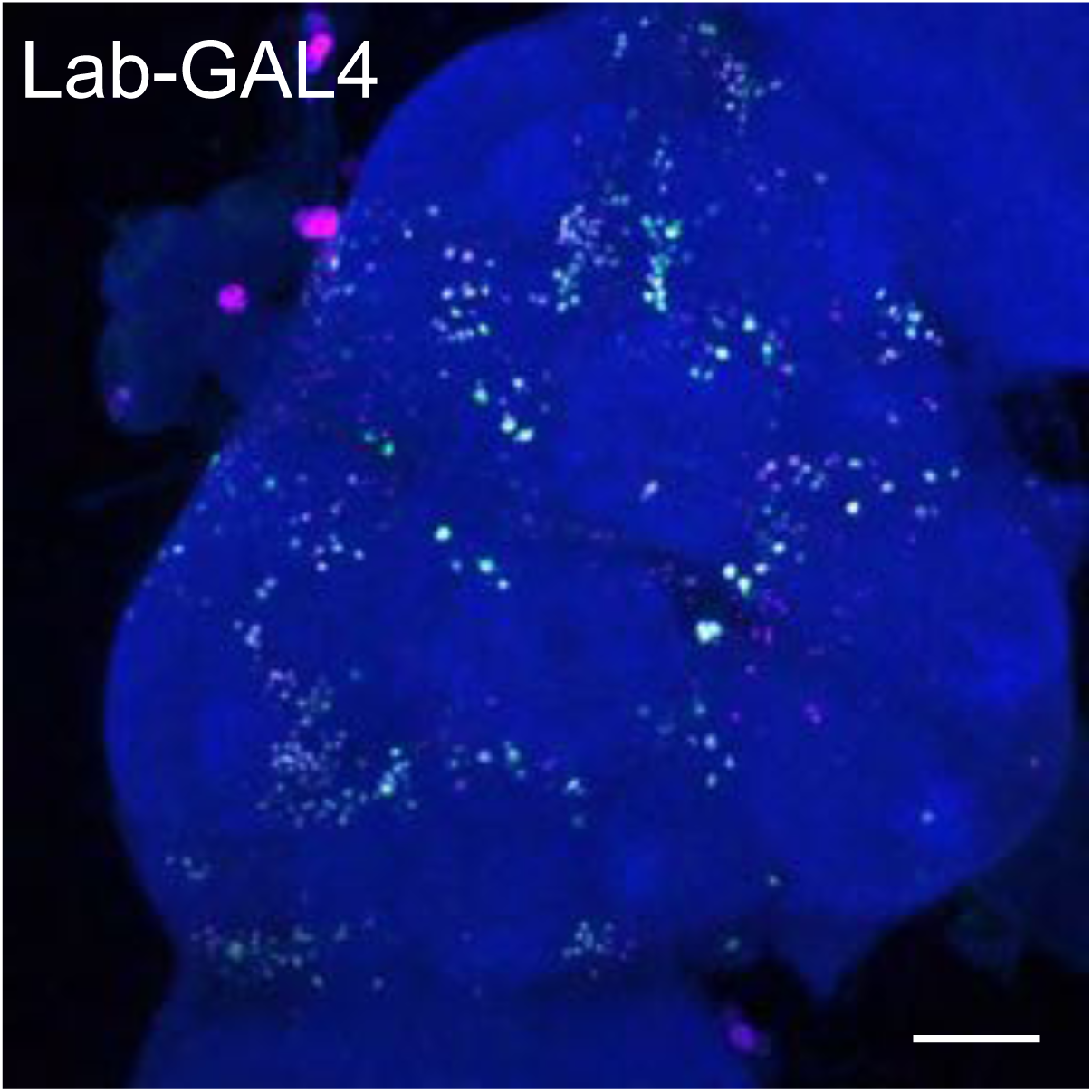
Expression of labial-GAL4 in the adult central brain. Labial-GAL4 driving UAS-nls-GCaMP6s (green) and UAS-His2Av-mRFP (fuchsia) in an adult female brain. Neuropil is labeled with nc82 (blue). Cell bodies throughout the central brain are labeled. Scale bar 50 μm.

**Figure 2—figure supplement 1.**
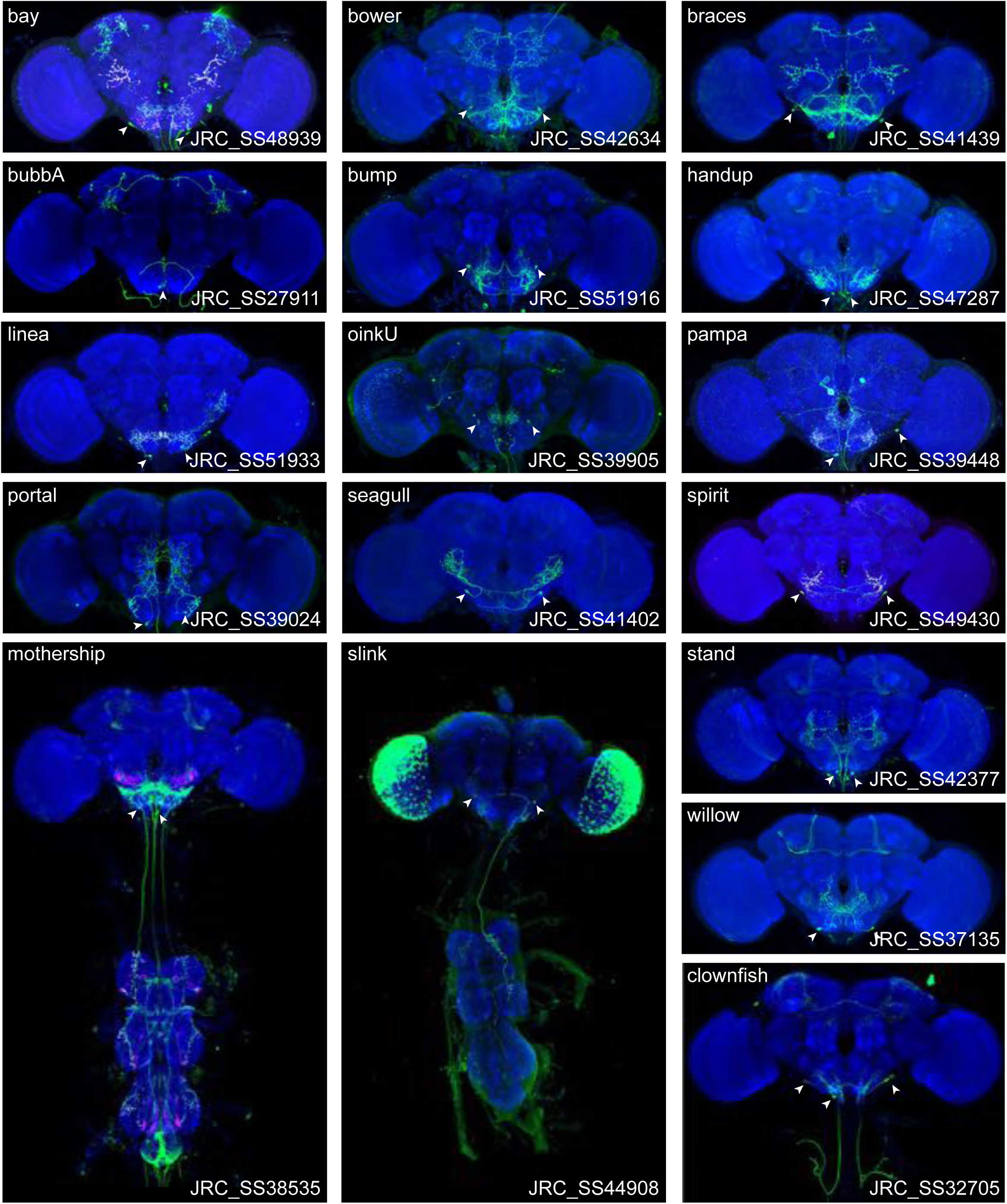
Expression patterns of split-GAL4 lines targeting neuronal cell types not included in NBLAST clustering. Neuropil morphology is shown with nc82 in blue. Expression pattern of the UAS reporter is shown in green. UAS-Synaptotagmin staining to indicate location of synaptic outputs is shown in magenta where available. The cell type covered is indicated in the top right of each panel, while the unique line identifier is indicated in the bottom left of each panel. Filled arrowheads indicate targeted cell type somas. See https://splitgal4.janelia.org/ for image data.

**Figure 2—figure supplement 2.**
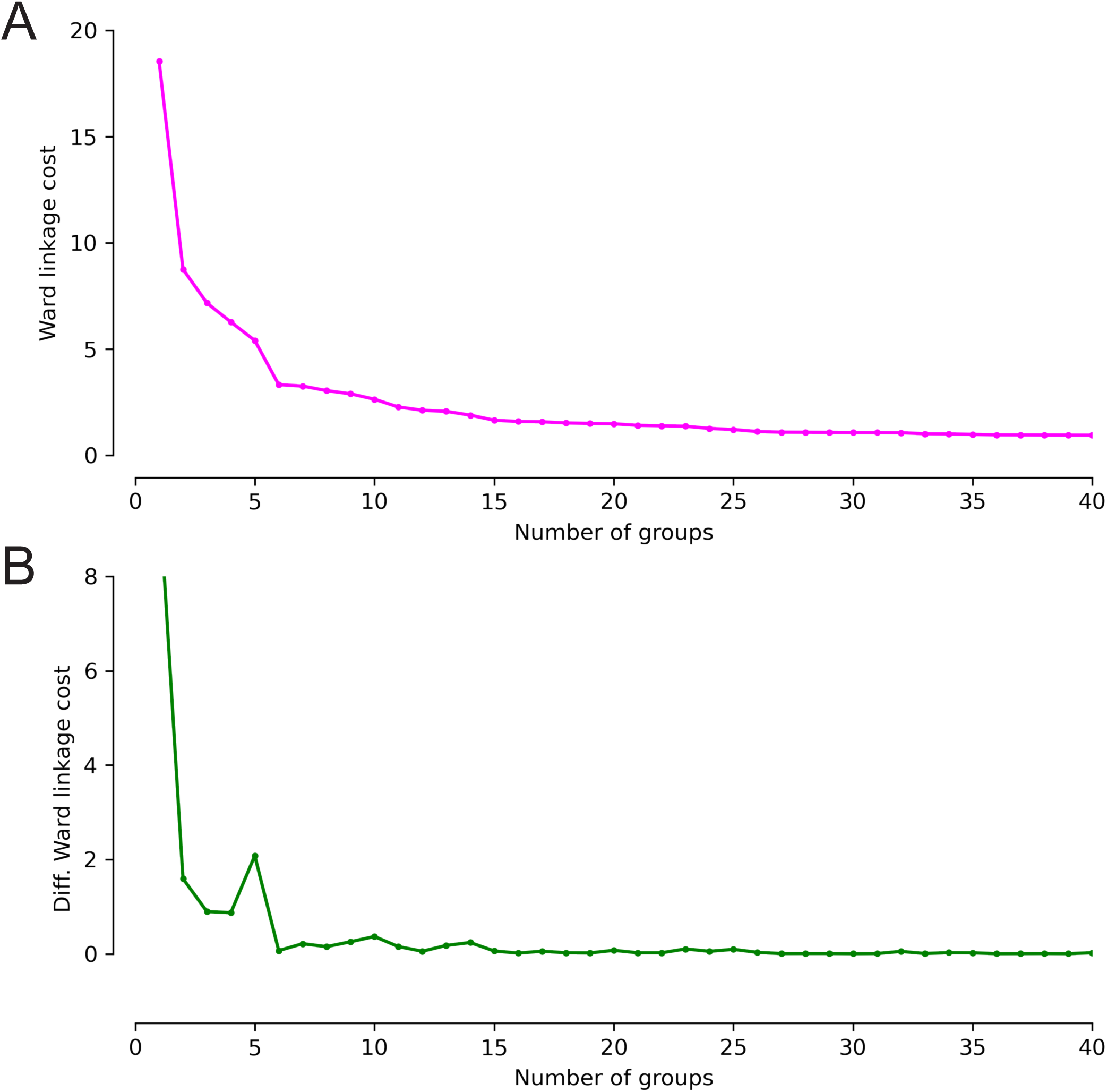
Ward’s joining cost and the differential of Ward’s joining cost for hierarchical clustering of SEZ neuronal cell types with NBLAST. (A) Ward’s joining cost for clustering into 0 to 40 groups (fuchsia). Ward’s joining cost declines sharply when clustering with 6 groups as compared to clustering with fewer than 6 groups. (B) Differential of Ward’s joining cost for clustering into 0 to 40 groups (green). The differential is high when clustering into 5 groups or fewer but does not decline notably after 6 groups is reached.

**Figure 3—figure supplement 1.**
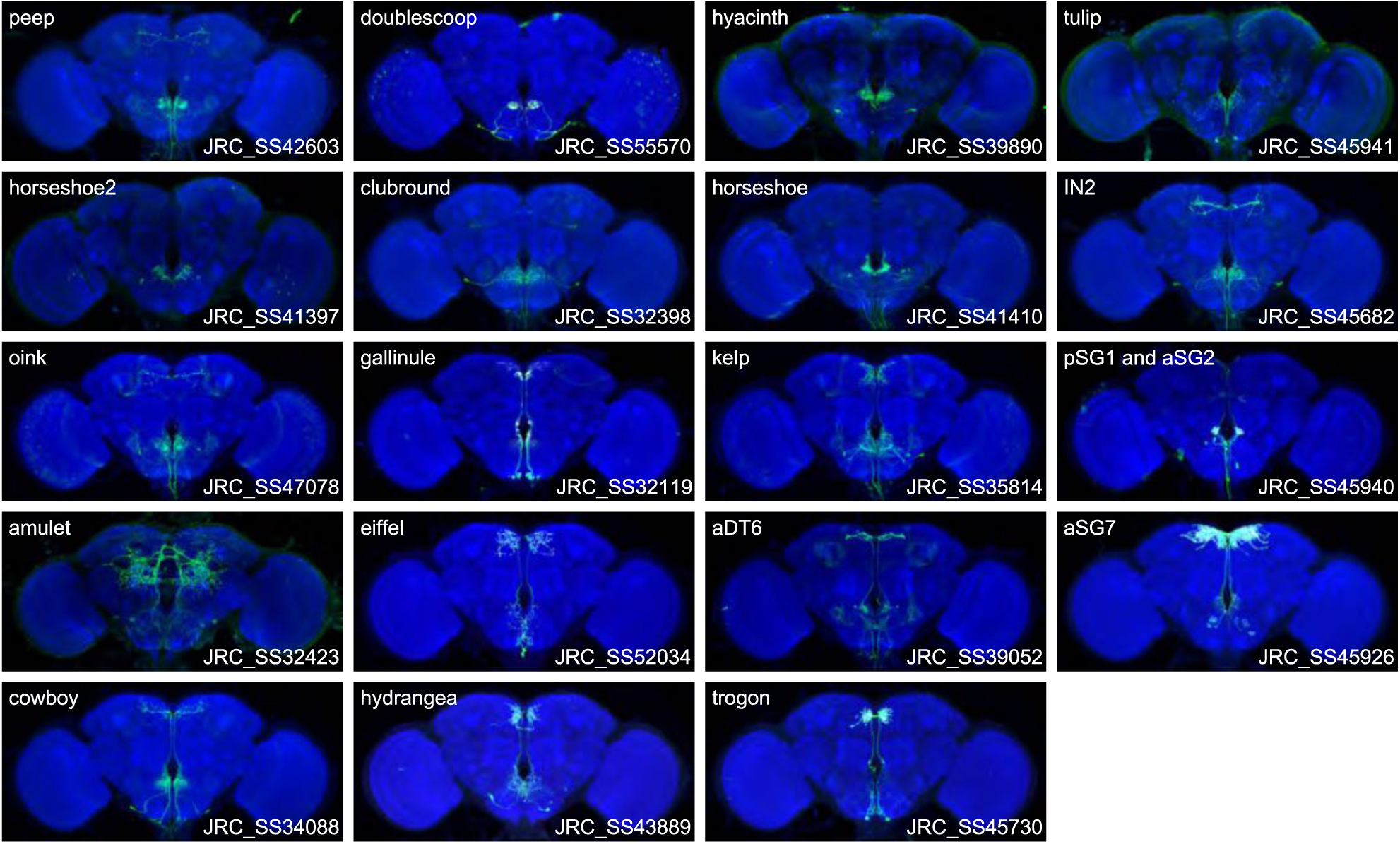
Expression patterns of the best split-GAL4 for each neuron type in Group 1. Neuropil morphology is shown with nc82 in blue throughout. Expression pattern of the UAS reporter is shown in green. UAS-Synaptotagmin staining indicates the location of synaptic outputs in magenta where available. The cell type covered is indicated in the top right of each panel, while the unique line identifier is indicated in the bottom left of each panel. See https://splitgal4.janelia.org/ for image data.

**Figure 4 – figure supplement 1.**
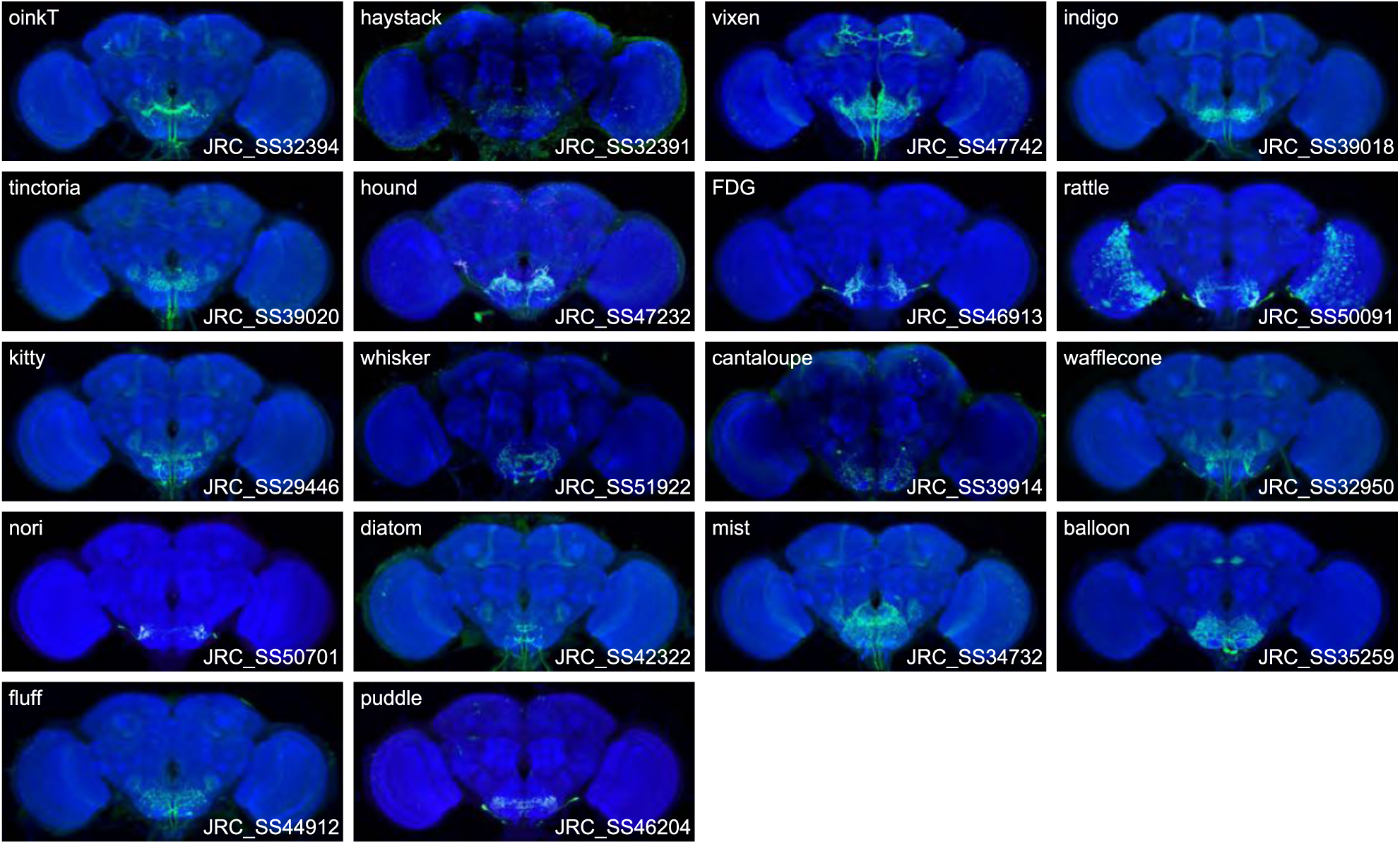
Expression patterns of the best split-GAL4 for each neuron type in Group 2. Neuropil morphology is shown with nc82 in blue throughout. Expression pattern of the UAS reporter is shown in green. UAS-Synaptotagmin staining indicates the location of synaptic outputs in magenta where available. The cell type covered is indicated in the top right of each panel, while the unique line identifier is indicated in the bottom left of each panel. See https://splitgal4.janelia.org/ for image data.

**Figure 4 – figure supplement 2.**
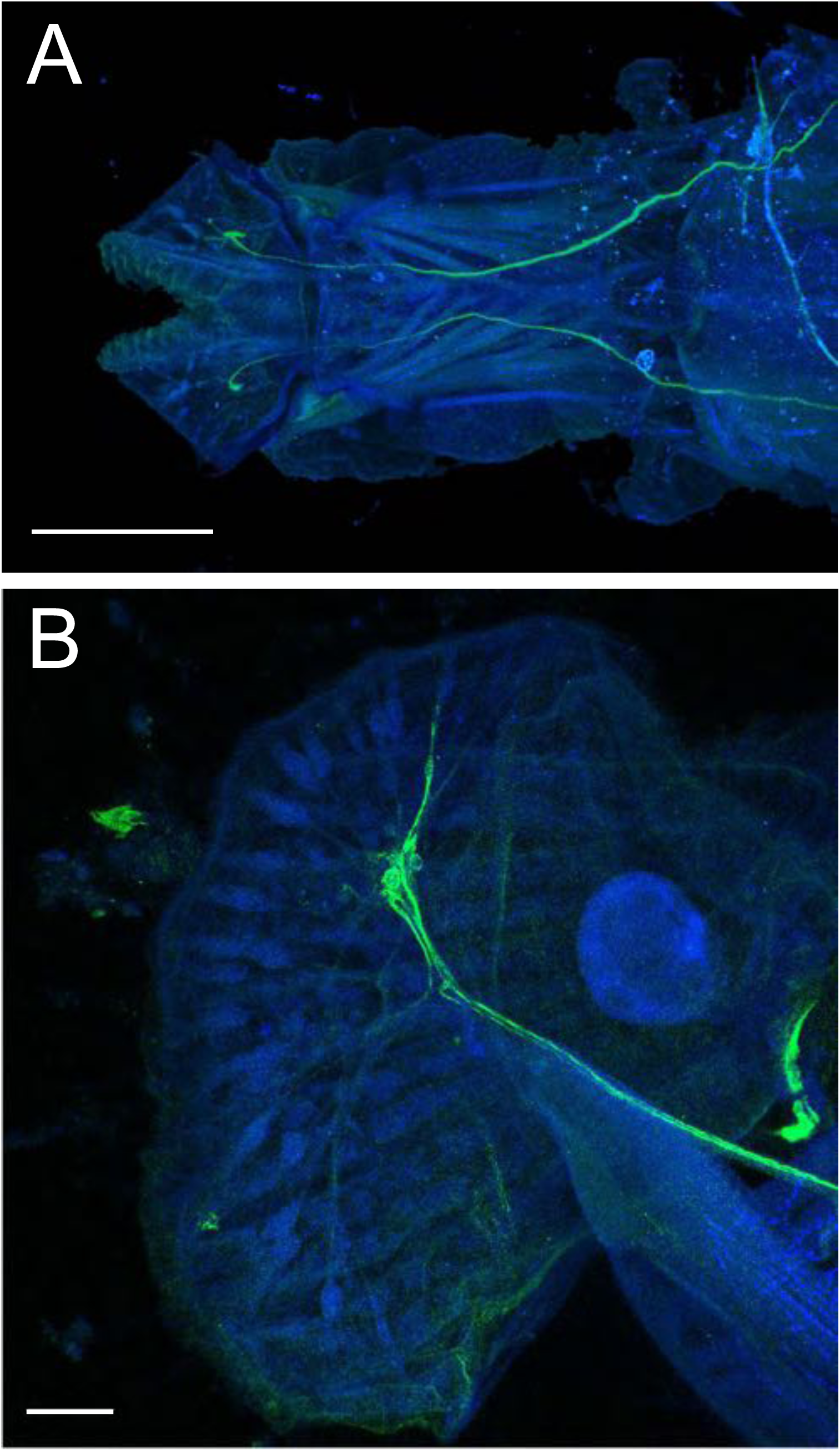
Diatom cell bodies and dendrites in the proboscis labellum. (A) Dorsal view of a whole mount proboscis showing the expression pattern of JRC_SS40945. The labellum (left) contains the cell bodies of diatom. The axons project through the labellar nerve into the SEZ. Scale bar is 100 μm. (B) Higher resolution lateral image of the proboscis labellum showing the expression pattern of JRC_SS40945. The dendrites of diatom project toward the dorsal surface of the labellum from the cell bodies. Scale bar is 20 μm. Both images show presynaptic sites impinging on proboscis muscles with nc82 in blue, while the expression of the UAS-CsChrimson reporter is shown in green.

**Figure 5 – figure supplement 1.**
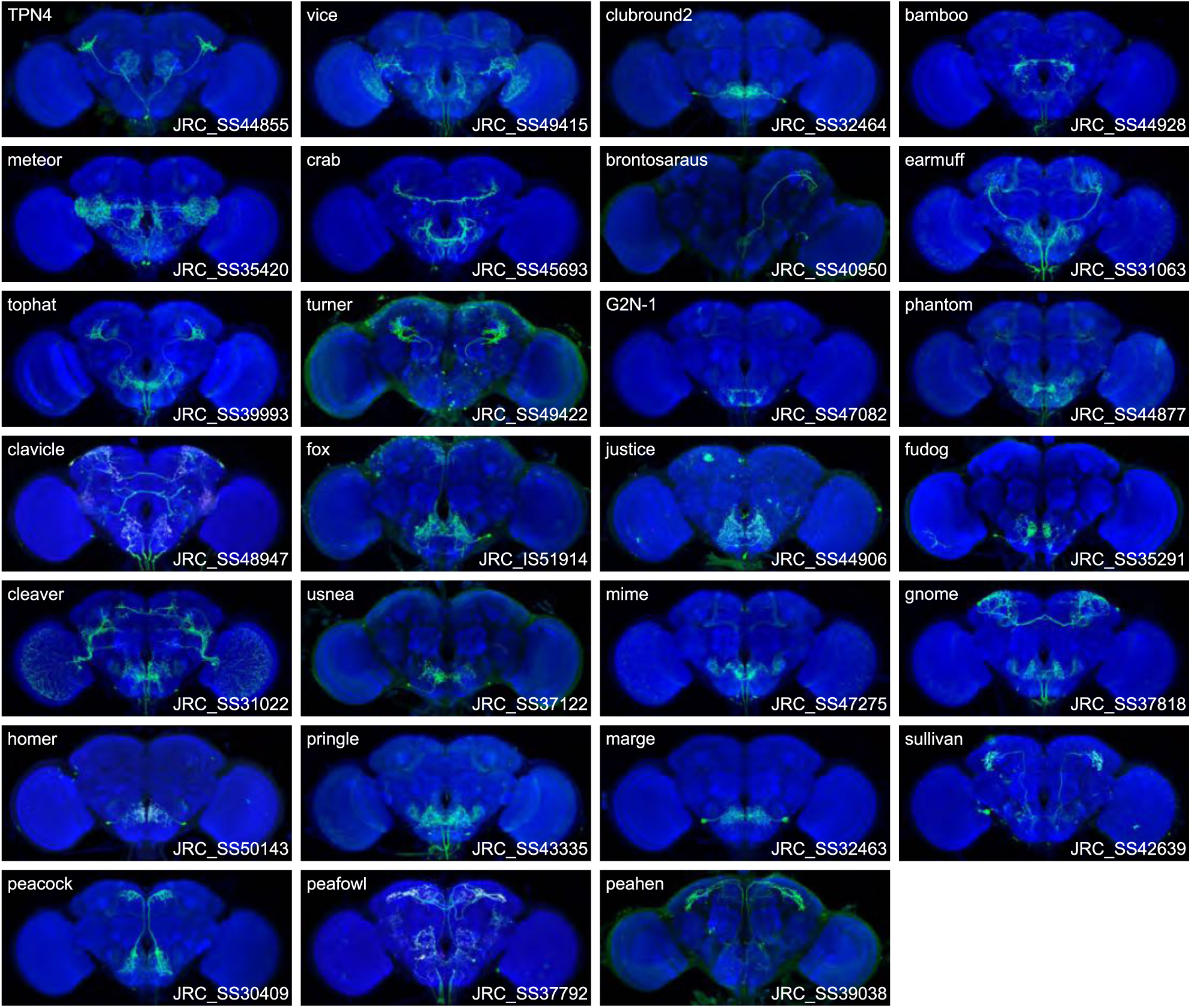
Expression patterns of the best split-GAL4 for each neuron type in Group 3. Neuropil morphology is shown with nc82 in blue throughout. Expression pattern of the UAS reporter is shown in green. UAS-Synaptotagmin staining indicates the location of synaptic outputs in magenta where available. The cell type covered is indicated in the top right of each panel, while the unique line identifier is indicated in the bottom left of each panel. See https://splitgal4.janelia.org/ for image data.

**Figure 6—figure supplement 1.**
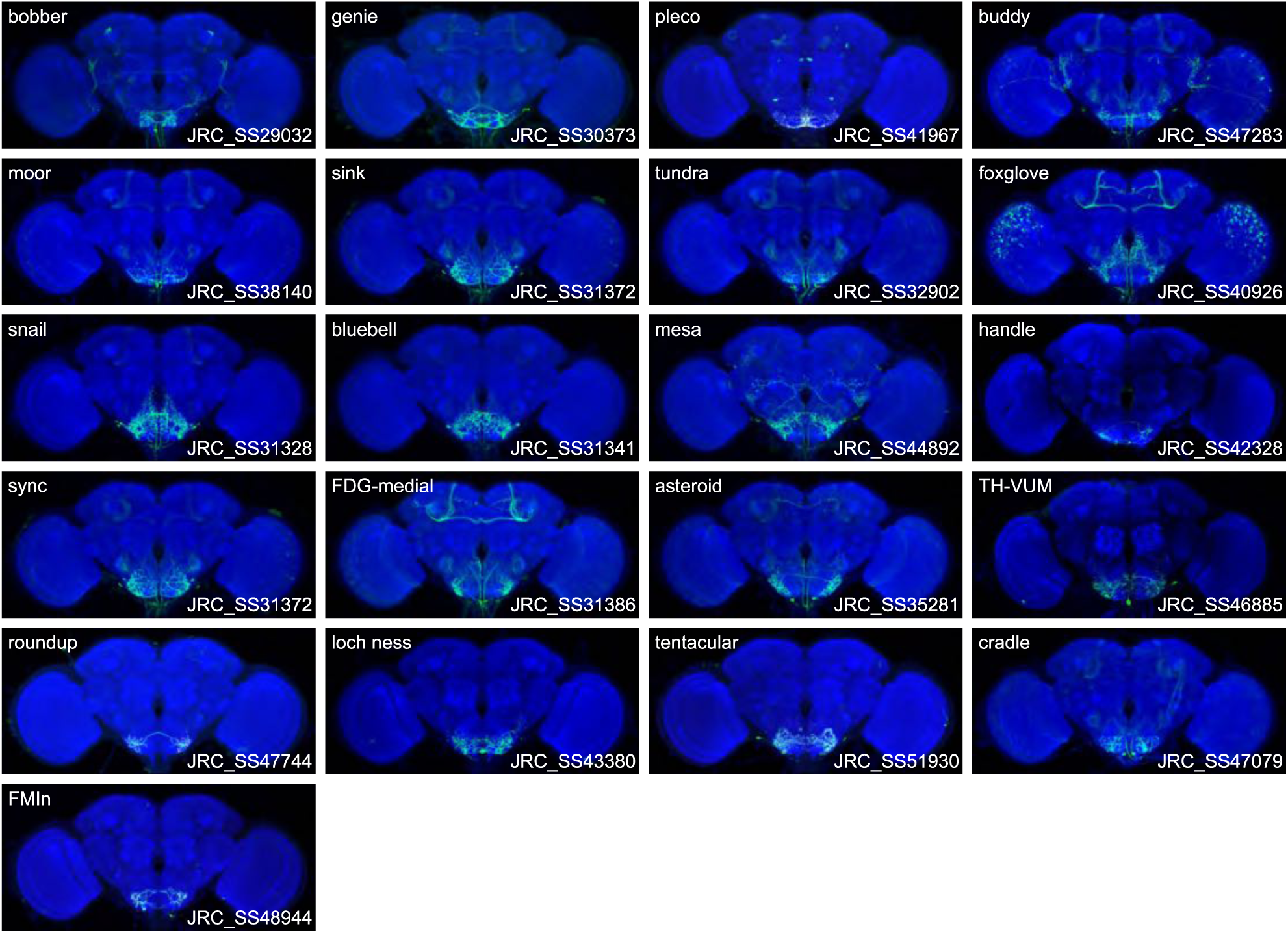
Expression patterns of the best split-GAL4 for each neuron type in Group 4. Neuropil morphology is shown with nc82 in blue throughout. Expression pattern of the UAS reporter is shown in green. UAS-Synaptotagmin staining indicates the location of synaptic outputs in magenta where available. The cell type covered is indicated in the top right of each panel, while the unique line identifier is indicated in the bottom left of each panel. See https://splitgal4.janelia.org/ for image data.

**Figure 6—figure supplement 2.**
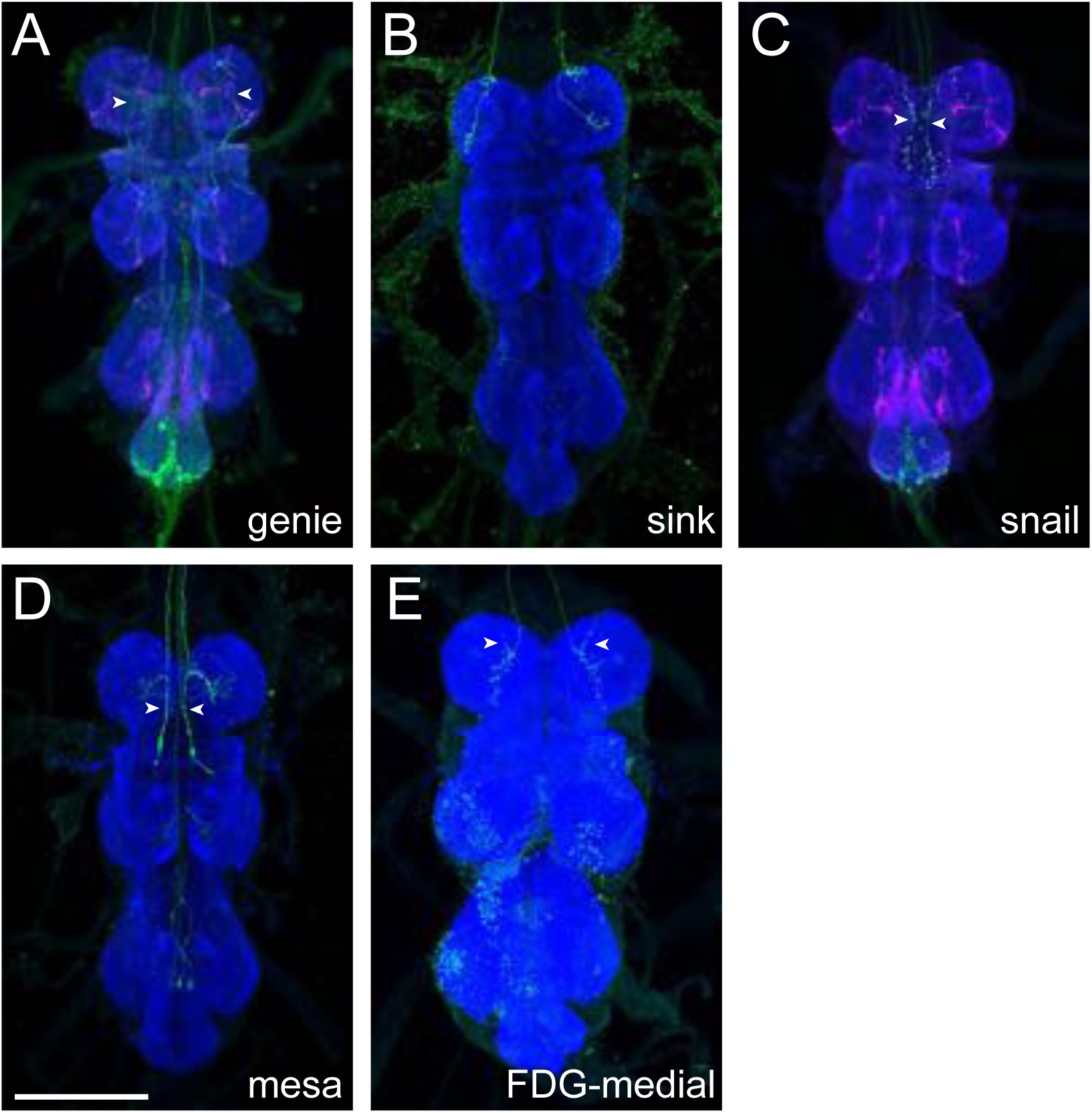
Axonal morphology of descending interneurons clustered into Group 4. (A) Genie axonal projections in the pattern of JRC_SS30373. (B) Sink axonal projections in the pattern of JRC_SS31372. (C) Snail cell type in the pattern of JRC_SS31328. (D) Mesa axonal projections in the pattern of JRC_SS31369. (E) FDG-medial axonal projections in the pattern of JRC_SS31386. Some non-specific background can be seen in the inferior VNC. Neuropil morphology is shown with nc82 in blue throughout. Expression pattern of the UAS reporter is shown in green. UAS-Synaptotagmin staining indicates the location of synaptic outputs in magenta where available (A and C), but note non-specific background throughout the VNC). White arrowheads denote the axons of the cell type of interest where other cell types are present. All images are unregistered VNCs. Scale bar 100 μm.

**Figure 7—figure supplement 1.**
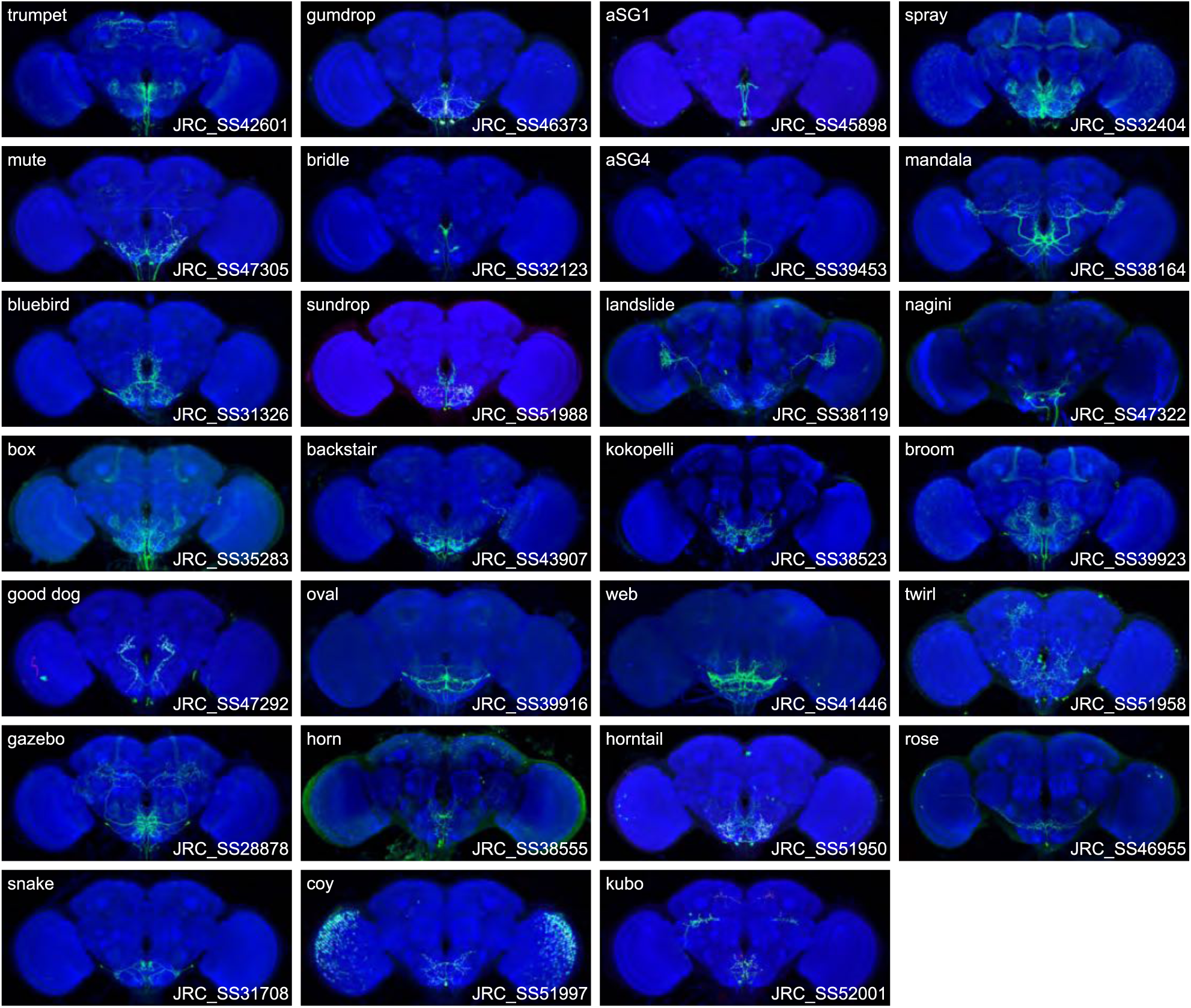
Expression patterns of the best split-GAL4 for each neuron type in Group 5. Neuropil morphology is shown with nc82 in blue throughout. Expression pattern of the UAS reporter is shown in green. UAS-Synaptotagmin staining indicates the location of synaptic outputs in magenta where available. The cell type covered is indicated in the top right of each panel, while the unique line identifier is indicated in the bottom left of each panel. See https://splitgal4.janelia.org/ for image data.

**Figure 7—figure supplement 2.**
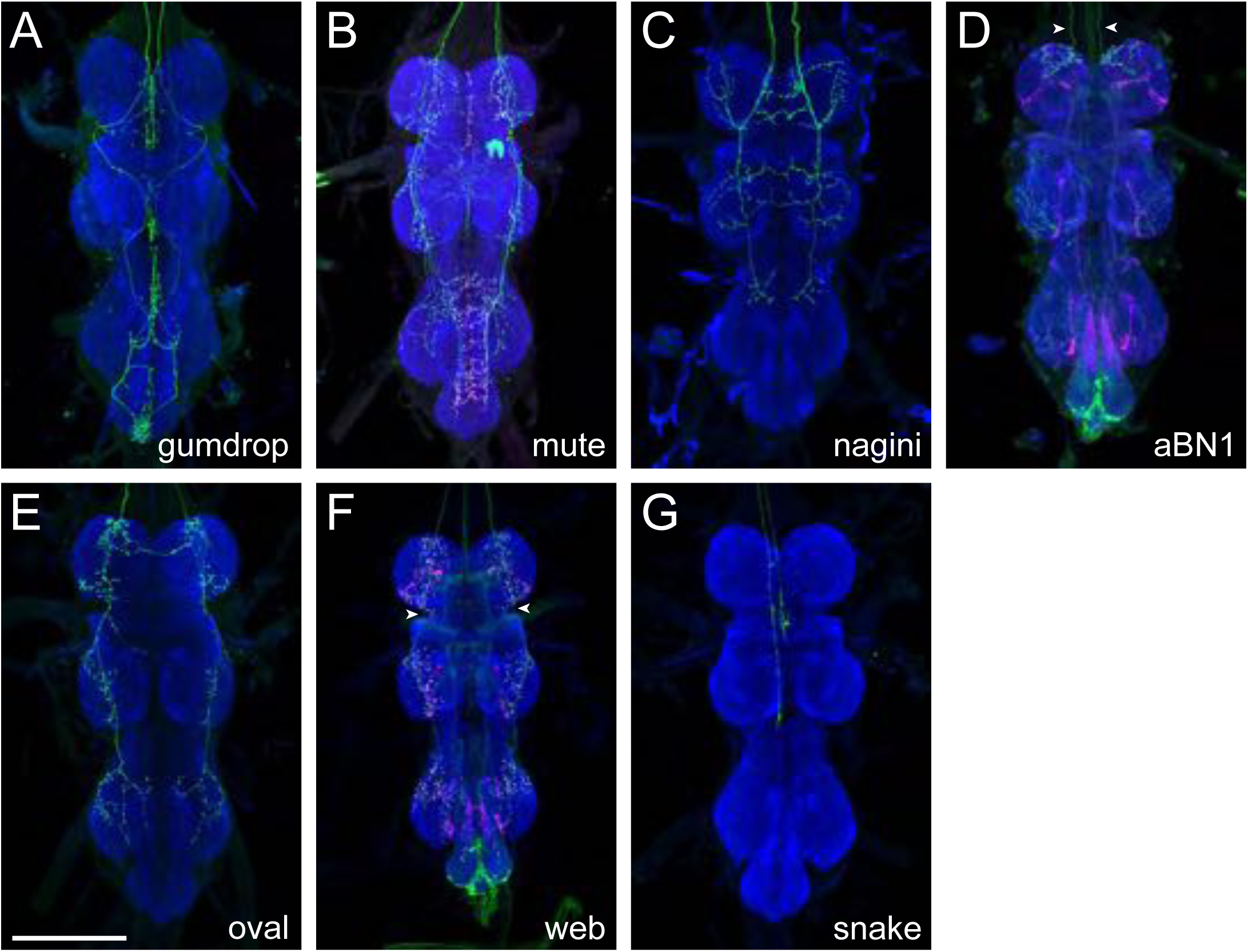
Axonal morphology of descending interneurons clustered into Group 5. (A) Gumdrop axonal projections in the pattern of JRC_SS46373. (B) Mute axonal projections in the pattern of JRC_SS47305. (C) Nagini axonal projections in the pattern of JRC_SS47322. (D) aBN1 axonal projections in the pattern of JRC_SS43907. (E) Oval axonal projections in the pattern of JRC_SS39916. (F) Web axonal projections in the pattern of JRC_SS41446. (G) Snake axonal projections in the pattern of JRC_SS31714. Neuropil morphology is shown with nc82 in blue throughout. Expression pattern of the UAS reporter is shown in green. UAS-Synaptotagmin staining indicates the location of synaptic outputs in magenta where available (B, D, and F, but note non-specific background throughout the VNC). White arrowheads denote the axons of the cell type of interest where other cell types are present. All images are unregistered VNCs. Scale bar 100 μm.

**Figure 8—figure supplement 1.**
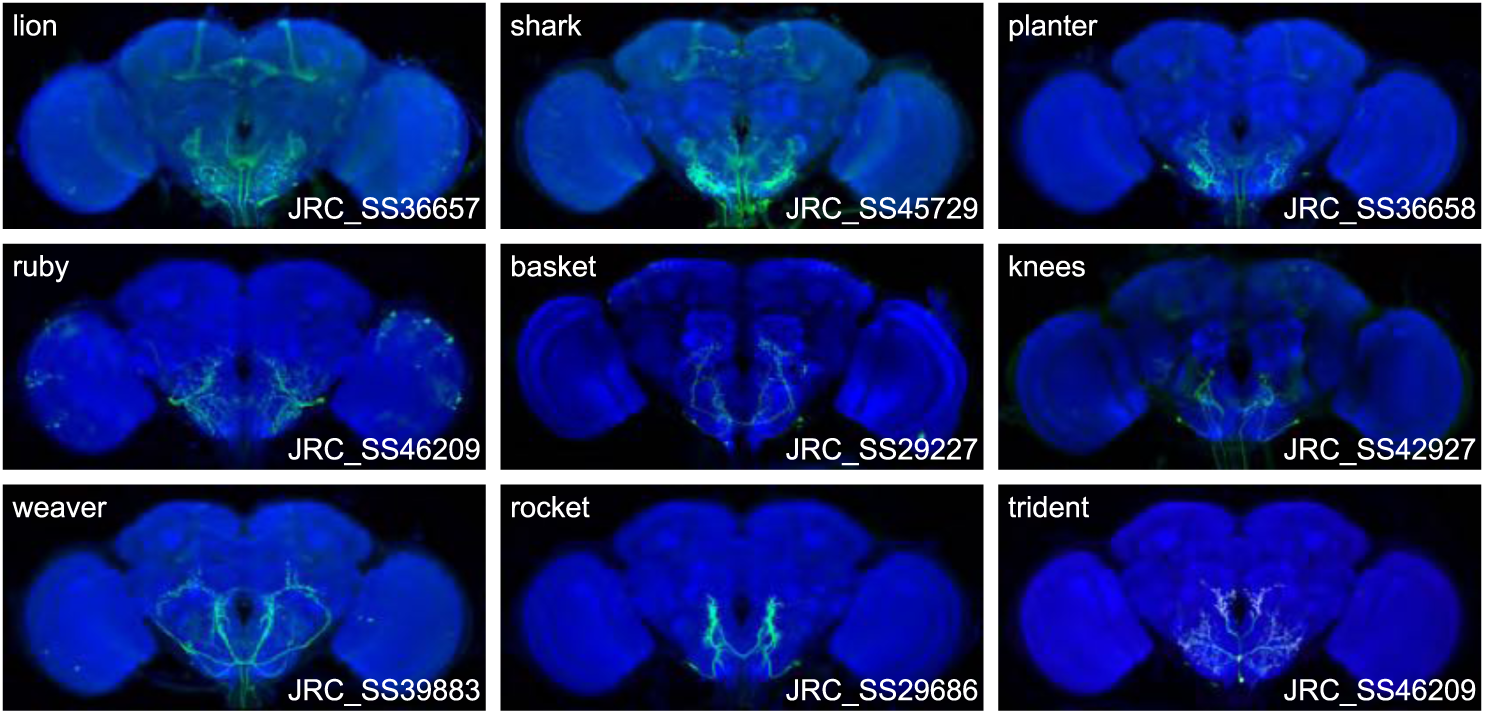
Expression patterns of the best split-GAL4 for each neuron type in Group 6. Neuropil morphology is shown with nc82 in blue throughout. Expression pattern of the UAS reporter is shown in green. UAS-Synaptotagmin staining indicates the location of synaptic outputs in magenta where available. The cell type covered is indicated in the top right of each panel, while the unique line identifier is indicated in the bottom left of each panel. See https://splitgal4.janelia.org/ for image data.

## References

Aso Y, Hattori D, Yu Y, Johnston RM, Iyer NA, Ngo T-TB, Dionne H, Abbott LF, Axel R, Tanimoto H, Rubin GM. 2014a. The neuronal architecture of the mushroom body provides a logic for associative learning. Elife 3:e04577. doi:10.7554/eLife.04577

Aso Y, Sitaraman D, Ichinose T, Kaun KR, Vogt K, Belliart-Guérin G, Plaçais P-Y, Robie AA, Yamagata N, Schnaitmann C, Rowell WJ, Johnston RM, Ngo T-TB, Chen N, Korff W, Nitabach MN, Heberlein U, Preat T, Branson KM, Tanimoto H, Rubin GM. 2014b. Mushroom body output neurons encode valence and guide memory-based action selection in Drosophila. eLife 3:e04580. doi:10.7554/eLife.04580

Bates AS, Manton JD, Jagannathan SR, Costa M, Schlegel P, Rohlfing T, Jefferis GS. 2020. The natverse, a versatile toolbox for combining and analysing neuroanatomical data. eLife 9:e53350. doi:10.7554/eLife.53350

Berg S, Kutra D, Kroeger T, Straehle CN, Kausler BX, Haubold C, Schiegg M, Ales J, Beier T, Rudy M, Eren K, Cervantes JI, Xu B, Beuttenmueller F, Wolny A, Zhang C, Koethe U, Hamprecht FA, Kreshuk A. 2019. ilastik: interactive machine learning for (bio)image analysis. Nature Methods 16:1226–1232. doi:10.1038/s41592-019-0582-9

Bidaye SS, Laturney M, Chang AK, Liu Y, Bockemühl T, Büschges A, Scott K. 2020. Two Brain Pathways Initiate Distinct Forward Walking Programs in Drosophila. Neuron 108:469–485.e8. doi:10.1016/j.neuron.2020.07.032

Bidaye SS, Machacek C, Wu Y, Dickson BJ. 2014. Neuronal Control of Drosophila Walking Direction. Science 344:97–101. doi:10.1126/science.1249964

Bogovic JA, Otsuna H, Heinrich L, Ito M, Jeter J, Meissner G, Nern A, Colonell J, Malkesman O, Ito K, Saalfeld S. 2020. An unbiased template of the Drosophila brain and ventral nerve cord. PLoS One 15. doi:10.1371/journal.pone.0236495

Boyan G, Reichert H, Hirth F. 2003. Commissure formation in the embryonic insect brain. Arthropod Structure & Development, Development of the Arthropod Nervous System: a Comparative and Evolutionary Approach 32:61–77. doi:10.1016/S1467-8039(03)00037-9

Braun E, Geurten B, Egelhaaf M. 2010. Identifying Prototypical Components in Behaviour Using Clustering Algorithms. PLoS One 5. doi:10.1371/journal.pone.0009361

Chiang A-S, Lin Chih-Yung, Chuang C-C, Chang H-M, Hsieh C-H, Yeh C-W, Shih C-T, Wu J-J, Wang G-T, Chen Y-C, Wu Cheng-Chi, Chen G-Y, Ching Y-T, Lee P-C, Lin Chih-Yang, Lin H-H, Wu Chia-Chou, Hsu H-W, Huang Y-A, Chen J-Y, Chiang H-J, Lu C-F, Ni R-F, Yeh C-Y, Hwang J-K. 2011. Three-Dimensional Reconstruction of Brain-wide Wiring Networks in Drosophila at Single-Cell Resolution. Current Biology 21:1–11. doi:10.1016/j.cub.2010.11.056

Costa M, Manton JD, Ostrovsky AD, Prohaska S, Jefferis GSXE. 2016. NBLAST: Rapid, Sensitive Comparison of Neuronal Structure and Construction of Neuron Family Databases. Neuron 91:293–311. doi:10.1016/j.neuron.2016.06.012

Court R, Namiki S, Armstrong JD, Börner J, Card G, Costa M, Dickinson M, Duch C, Korff W, Mann R, Merritt D, Murphey RK, Seeds AM, Shirangi T, Simpson JH, Truman JW, Tuthill JC, Williams DW, Shepherd D. 2020. A Systematic Nomenclature for the Drosophila Ventral Nerve Cord. Neuron 107:1071–1079.e2. doi:10.1016/j.neuron.2020.08.005

Davie K, Janssens J, Koldere D, De Waegeneer M, Pech U, Kreft Ł, Aibar S, Makhzami S, Christiaens V, Bravo González-Blas C, Poovathingal S, Hulselmans G, Spanier KI, Moerman T, Vanspauwen B, Geurs S, Voet T, Lammertyn J, Thienpont B, Liu S, Konstantinides N, Fiers M, Verstreken P, Aerts S. 2018. A Single-Cell Transcriptome Atlas of the Aging Drosophila Brain. Cell 174:982–998.e20. doi:10.1016/j.cell.2018.05.057

Dionne H, Hibbard KL, Cavallaro A, Kao J-C, Rubin GM. 2018. Genetic Reagents for Making Split-GAL4 Lines in Drosophila. Genetics 209:31–35. doi:10.1534/genetics.118.300682

Dolan M-J, Frechter S, Bates AS, Dan C, Huoviala P, Roberts RJ, Schlegel P, Dhawan S, Tabano R, Dionne H, Christoforou C, Close K, Sutcliffe B, Giuliani B, Li F, Costa M, Ihrke G, Meissner GW, Bock DD, Aso Y, Rubin GM, Jefferis GS. 2019. Neurogenetic dissection of the Drosophila lateral horn reveals major outputs, diverse behavioural functions, and interactions with the mushroom body. eLife 8:e43079. doi:10.7554/eLife.43079

Dolan M-J, Luan H, Shropshire WC, Sutcliffe B, Cocanougher B, Scott RL, Frechter S, Zlatic M, Jefferis GSXE, White BH. 2017. Facilitating Neuron-Specific Genetic Manipulations in Drosophila melanogaster Using a Split GAL4 Repressor. Genetics 206:775–784. doi:10.1534/genetics.116.199687

Dorkenwald S, McKellar C, Macrina T, Kemnitz N, Lee K, Lu R, Wu J, Popovych S, Mitchell E, Nehoran B, Jia Z, Bae JA, Mu S, Ih D, Castro M, Ogedengbe O, Halageri A, Ashwood Z, Zung J, Brittain D, Collman F, Schneider-Mizell C, Jordan C, Silversmith W, Baker C, Deutsch D, Encarnacion-Rivera L, Kumar S, Burke A, Gager J, Hebditch J, Koolman S, Moore M, Morejohn S, Silverman B, Willie K, Willie R, Yu S, Murthy M, Seung HS. 2020. FlyWire: Online community for whole-brain connectomics. bioRxiv 2020.08.30.274225. doi:10.1101/2020.08.30.274225

Emery G, Hutterer A, Berdnik D, Mayer B, Wirtz-Peitz F, Gaitan MG, Knoblich JA. 2005. Asymmetric Rab11 Endosomes Regulate Delta Recycling and Specify Cell Fate in the Drosophila Nervous System. Cell 122:763–773. doi:10.1016/j.cell.2005.08.017

Flood TF, Iguchi S, Gorczyca M, White B, Ito K, Yoshihara M. 2013. A single pair of interneurons commands the Drosophila feeding motor program. Nature 499:83–87. doi:10.1038/nature12208

Gordon MD, Scott K. 2009. Motor control in a Drosophila taste circuit. Neuron 61:373–384. doi:10.1016/j.neuron.2008.12.033

Haberkern H, Basnak MA, Ahanonu B, Schauder D, Cohen JD, Bolstad M, Bruns C, Jayaraman V. 2019. Visually Guided Behavior and Optogenetically Induced Learning in Head-Fixed Flies Exploring a Virtual Landscape. Current Biology 29:1647–1659.e8. doi:10.1016/j.cub.2019.04.033

Hampel S, Franconville R, Simpson JH, Seeds AM. 2015. A neural command circuit for grooming movement control. eLife 4:e08758. doi:10.7554/eLife.08758

Hampel S, McKellar CE, Simpson JH, Seeds AM. 2017. Simultaneous activation of parallel sensory pathways promotes a grooming sequence in Drosophila. eLife. doi:10.7554/eLife.28804

Harris DT, Kallman BR, Mullaney BC, Scott K. 2015. Representations of Taste Modality in the Drosophila Brain. Neuron 86:1449–1460. doi:10.1016/j.neuron.2015.05.026

Hartenstein V, Omoto JJ, Ngo KT, Wong D, Kuert PA, Reichert H, Lovick JK, Younossi-Hartenstein A. 2018. Structure and development of the subesophageal zone of the Drosophila brain. I. Segmental architecture, compartmentalization, and lineage anatomy. J Comp Neurol 526:6–32. doi:10.1002/cne.24287

Hirth F, Hartmann B, Reichert H. 1998. Homeotic gene action in embryonic brain development of Drosophila. Development 125:1579–1589.

Hirth F, Loop T, Egger B, Miller DF, Kaufman TC, Reichert H. 2001. Functional equivalence of Hox gene products in the specification of the tritocerebrum during embryonic brain development of Drosophila. Development 128:4781–4788.

Huang L-K, Wang M-JJ. 1995. Image thresholding by minimizing the measures of fuzziness. Pattern Recognit. doi:10.1016/0031-3203(94)E0043-K

Ito K, Shinomiya K, Ito M, Armstrong JD, Boyan G, Hartenstein V, Harzsch S, Heisenberg M, Homberg U, Jenett A, Keshishian H, Restifo LL, Rössler W, Simpson JH, Strausfeld NJ, Strauss R, Vosshall LB, Insect Brain Name Working Group. 2014. A systematic nomenclature for the insect brain. Neuron 81:755–765. doi:10.1016/j.neuron.2013.12.017

Jenett A, Rubin GM, Ngo T-TB, Shepherd D, Murphy C, Dionne H, Pfeiffer BD, Cavallaro A, Hall D, Jeter J, Iyer N, Fetter D, Hausenfluck JH, Peng H, Trautman ET, Svirskas RR, Myers EW, Iwinski ZR, Aso Y, DePasquale GM, Enos A, Hulamm P, Lam SCB, Li H-H, Laverty TR, Long F, Qu L, Murphy SD, Rokicki K, Safford T, Shaw K, Simpson JH, Sowell A, Tae S, Yu Y, Zugates CT. 2012. A GAL4-driver line resource for Drosophila neurobiology. Cell Rep 2:991–1001. doi:10.1016/j.celrep.2012.09.011

Jourjine N, Mullaney BC, Mann K, Scott K. 2016. Coupled Sensing of Hunger and Thirst Signals Balances Sugar and Water Consumption. Cell 166:855–866. doi:10.1016/j.cell.2016.06.046

Kendroud S, Bohra AA, Kuert PA, Nguyen B, Guillermin O, Sprecher SG, Reichert H, VijayRaghavan K, Hartenstein V. 2018. Structure and development of the subesophageal zone of the Drosophila brain. II. Sensory compartments. Journal of Comparative Neurology 526:33–58. doi:https://doi.org/10.1002/cne.24316

Kim H, Kirkhart C, Scott K. 2017. Long-range projection neurons in the taste circuit of Drosophila. eLife 6:e23386. doi:10.7554/eLife.23386

Klapoetke NC, Murata Y, Kim SS, Pulver SR, Birdsey-Benson A, Cho YK, Morimoto TK, Chuong AS, Carpenter EJ, Tian Z, Wang J, Xie Y, Yan Z, Zhang Y, Chow BY, Surek B, Melkonian M, Jayaraman V, Constantine-Paton M, Wong GK-S, Boyden ES. 2014. Independent optical excitation of distinct neural populations. Nature Methods 11:338–346. doi:10.1038/nmeth.2836

Kuert PA, Hartenstein V, Bello BC, Lovick JK, Reichert H. 2014. Neuroblast lineage identification and lineage-specific Hox gene action during postembryonic development of the subesophageal ganglion in the Drosophila central brain. Developmental Biology 390:102–115. doi:10.1016/j.ydbio.2014.03.021

Kumar R, Chotaliya M, Vuppala S, Auradkar A, Palasamudrum K, Joshi R. 2015. Role of Homothorax in region specific regulation of Deformed in embryonic neuroblasts. Mech Dev 138:190–197. doi:10.1016/j.mod.2015.09.003

Lee T-C, Kashyap RL, Chu C-N. 1994. Building skeleton models via 3-D medial surface/axis thinning algorithms. CVGIP: Graph Models Image Process 56:462–478. doi:10.1006/cgip.1994.1042

Ling F, Dahanukar A, Weiss LA, Kwon JY, Carlson JR. 2014. The molecular and cellular basis of taste coding in the legs of Drosophila. J Neurosci 34:7148–7164. doi:10.1523/JNEUROSCI.0649-14.2014

Liu, Tianxiao. 2012. Anatomical and functional dissection of fruitless positive courtship circuit in Drosophila melanogaster. Vienna, Austria: University of Vienna.

Luan H, Peabody NC, Vinson CR, White BH. 2006. Refined spatial manipulation of neuronal function by combinatorial restriction of transgene expression. Neuron 52:425–436. doi:10.1016/j.neuron.2006.08.028

Mann K, Gordon MD, Scott K. 2013. A pair of interneurons influences the choice between feeding and locomotion in Drosophila. Neuron 79:754–765. doi:10.1016/j.neuron.2013.06.018

Manzo A, Silies M, Gohl DM, Scott K. 2012. Motor neurons controlling fluid ingestion in Drosophila. PNAS 109:6307–6312. doi:10.1073/pnas.1120305109

Marella S, Fischler W, Kong P, Asgarian S, Rueckert E, Scott K. 2006. Imaging taste responses in the fly brain reveals a functional map of taste category and behavior. Neuron 49:285–295. doi:10.1016/j.neuron.2005.11.037

Marella S, Mann K, Scott K. 2012. Dopaminergic Modulation of Sucrose Acceptance Behavior in Drosophila. Neuron 73:941–950. doi:10.1016/j.neuron.2011.12.032

McKellar CE, Siwanowicz I, Dickson BJ, Simpson JH. 2020. Controlling motor neurons of every muscle for fly proboscis reaching. eLife 9:e54978. doi:10.7554/eLife.54978

Meissner GW, Dorman Z, Nern A, Forster K, Gibney T, Jeter J, Johnson L, He Y, Lee K, Melton B, Yarbrough B, Clements J, Goina C, Otsuna H, Rokicki K, Svirskas RR, Aso Y, Card GM, Dickson BJ, Ehrhardt E, Goldammer J, Ito M, Korff W, Minegishi R, Namiki S, Rubin GM, Sterne G, Wolff T, Malkesman O, Team FP. 2020. An image resource of subdivided Drosophila GAL4-driver expression patterns for neuron-level searches. bioRxiv 2020.05.29.080473. doi:10.1101/2020.05.29.080473

Miyazaki T, Lin T-Y, Ito K, Lee C-H, Stopfer M. 2015. A gustatory second-order neuron that connects sucrose-sensitive primary neurons and a distinct region of the gnathal ganglion in the Drosophila brain. J Neurogenet 29:144–155. doi:10.3109/01677063.2015.1054993

Namiki S, Dickinson MH, Wong AM, Korff W, Card GM. 2018. The functional organization of descending sensory-motor pathways in Drosophila. eLife 7:e34272. doi:10.7554/eLife.34272

Nern A, Pfeiffer BD, Rubin GM. 2015. Optimized tools for multicolor stochastic labeling reveal diverse stereotyped cell arrangements in the fly visual system. PNAS 112:E2967–E2976. doi:10.1073/pnas.1506763112

Otsuna H, Ito M, Kawase T. 2018. Color depth MIP mask search: a new tool to expedite Split-GAL4 creation. bioRxiv 318006. doi:10.1101/318006

Pandey R, Heidmann S, Lehner CF. 2005. Epithelial re-organization and dynamics of progression through mitosis in Drosophila separase complex mutants. J Cell Sci 118:733–742. doi:10.1242/jcs.01663

Pfeiffer BD, Jenett A, Hammonds AS, Ngo T-TB, Misra S, Murphy C, Scully A, Carlson JW, Wan KH, Laverty TR, Mungall C, Svirskas R, Kadonaga JT, Doe CQ, Eisen MB, Celniker SE, Rubin GM. 2008. Tools for neuroanatomy and neurogenetics in Drosophila. Proc Natl Acad Sci U S A 105:9715–9720. doi:10.1073/pnas.0803697105

Pfeiffer K, Homberg U. 2014. Organization and Functional Roles of the Central Complex in the Insect Brain. Annual Review of Entomology 59:165–184. doi:10.1146/annurev-ento-011613-162031

Preibisch S, Saalfeld S, Tomancak P. 2009. Globally optimal stitching of tiled 3D microscopic image acquisitions. Bioinformatics 25:1463–1465. doi:10.1093/bioinformatics/btp184

Riabinina O, Vernon SW, Dickson BJ, Baines RA. 2019. Split-QF System for Fine-Tuned Transgene Expression in Drosophila. Genetics 212:53–63. doi:10.1534/genetics.119.302034

Robie AA, Hirokawa J, Edwards AW, Umayam LA, Lee A, Phillips ML, Card GM, Korff W, Rubin GM, Simpson JH, Reiser MB, Branson K. 2017. Mapping the Neural Substrates of Behavior. Cell 170:393–406.e28. doi:10.1016/j.cell.2017.06.032

Robinson IM, Ranjan R, Schwarz TL. 2002. Synaptotagmins I and IV promote transmitter release independently of Ca 2+ binding in the C 2 A domain. Nature 418:336–340. doi:10.1038/nature00915

Rokicki, Konrad, Bruns, Christopher M., Goina, Cristian, Schauder, David, Olbris, Donald J., Trautman, Eric T., Svirskas, Rob, Clements, Jody, Ackerman, David, Kazimiers, Antje, Foster, Leslie L., Dolafi, Tom, Bolstad, Mark, Otsuna, Hideo, Yu, Yang, Safford, Todd, Murphy, Sean D. 2019. Janelia Workstation Codebase. Janlia Research Campus.

Scheffer LK, Xu CS, Januszewski M, Lu Z, Takemura Shin-ya, Hayworth KJ, Huang GB, Shinomiya K, Maitlin-Shepard J, Berg S, Clements J, Hubbard PM, Katz WT, Umayam L, Zhao T, Ackerman D, Blakely T, Bogovic J, Dolafi T, Kainmueller D, Kawase T, Khairy KA, Leavitt L, Li PH, Lindsey L, Neubarth N, Olbris DJ, Otsuna H, Trautman ET, Ito M, Bates AS, Goldammer J, Wolff T, Svirskas R, Schlegel P, Neace E, Knecht CJ, Alvarado CX, Bailey DA, Ballinger S, Borycz JA, Canino BS, Cheatham N, Cook M, Dreher M, Duclos O, Eubanks B, Fairbanks K, Finley S, Forknall N, Francis A, Hopkins GP, Joyce EM, Kim S, Kirk NA, Kovalyak J, Lauchie SA, Lohff A, Maldonado C, Manley EA, McLin S, Mooney C, Ndama M, Ogundeyi O, Okeoma N, Ordish C, Padilla N, Patrick CM, Paterson T, Phillips EE, Phillips EM, Rampally N, Ribeiro C, Robertson MK, Rymer JT, Ryan SM, Sammons M, Scott AK, Scott AL, Shinomiya A, Smith C, Smith K, Smith NL, Sobeski MA, Suleiman A, Swift J, Takemura Satoko, Talebi I, Tarnogorska D, Tenshaw E, Tokhi T, Walsh JJ, Yang T, Horne JA, Li F, Parekh R, Rivlin PK, Jayaraman V, Costa M, Jefferis GS, Ito K, Saalfeld S, George R, Meinertzhagen IA, Rubin GM, Hess HF, Jain V, Plaza SM. 2020. A connectome and analysis of the adult Drosophila central brain. eLife 9:e57443. doi:10.7554/eLife.57443

Schindelin J, Arganda-Carreras I, Frise E, Kaynig V, Longair M, Pietzsch T, Preibisch S, Rueden C, Saalfeld S, Schmid B, Tinevez J-Y, White DJ, Hartenstein V, Eliceiri K, Tomancak P, Cardona A. 2012. Fiji: an open-source platform for biological-image analysis. Nature Methods 9:676–682. doi:10.1038/nmeth.2019

Shirangi TR, Wong AM, Truman JW, Stern DL. 2016. Doublesex Regulates the Connectivity of a Neural Circuit Controlling Drosophila Male Courtship Song. Dev Cell 37:533–544. doi:10.1016/j.devcel.2016.05.012

Simpson JH. 2016. Rationally subdividing the fly nervous system with versatile expression reagents. Journal of Neurogenetics 30:185–194. doi:10.1080/01677063.2016.1248761

Snell NJ, Fisher JD, Hartmann GG, Talay M, Barnea G. 2020. Distributed Representation of Taste Quality by Second-Order Gustatory Neurons in Drosophila. bioRxiv 2020.11.10.377382. doi:10.1101/2020.11.10.377382

Stocker RF, Lienhard MC, Borst A, Fischbach KF. 1990. Neuronal architecture of the antennal lobe in Drosophila melanogaster. Cell Tissue Res 262:9–34. doi:10.1007/BF00327741

Tastekin I, Riedl J, Schilling-Kurz V, Gomez-Marin A, Truman JW, Louis M. 2015. Role of the subesophageal zone in sensorimotor control of orientation in Drosophila larva. Curr Biol 25:1448–1460. doi:10.1016/j.cub.2015.04.016

Thistle R, Cameron P, Ghorayshi A, Dennison L, Scott K. 2012. Contact chemoreceptors mediate male-male repulsion and male-female attraction during Drosophila courtship. Cell 149:1140–1151. doi:10.1016/j.cell.2012.03.045

Thorne N, Chromey C, Bray S, Amrein H. 2004. Taste perception and coding in Drosophila. Curr Biol 14:1065–1079. doi:10.1016/j.cub.2004.05.019

Ting C-Y, Gu S, Guttikonda S, Lin T-Y, White BH, Lee C-H. 2011. Focusing Transgene Expression in Drosophila by Coupling Gal4 With a Novel Split-LexA Expression System. Genetics 188:229–233. doi:10.1534/genetics.110.126193

Tirian L, Dickson BJ. 2017. The VT GAL4, LexA, and split-GAL4 driver line collections for targeted expression in the Drosophila nervous system. bioRxiv 198648. doi:10.1101/198648

Toda H, Zhao X, Dickson BJ. 2012. The Drosophila female aphrodisiac pheromone activates ppk23(+) sensory neurons to elicit male courtship behavior. Cell Rep 1:599–607. doi:10.1016/j.celrep.2012.05.007

Wang Z, Singhvi A, Kong P, Scott K. 2004. Taste representations in the Drosophila brain. Cell 117:981–991. doi:10.1016/j.cell.2004.06.011

Weiss LA, Dahanukar A, Kwon JY, Banerjee D, Carlson JR. 2011. The molecular and cellular basis of bitter taste in Drosophila. Neuron 69:258–272. doi:10.1016/j.neuron.2011.01.001

Wolff C, Tinevez J-Y, Pietzsch T, Stamataki E, Harich B, Guignard L, Preibisch S, Shorte S, Keller PJ, Tomancak P, Pavlopoulos A. 2018. Multi-view light-sheet imaging and tracking with the MaMuT software reveals the cell lineage of a direct developing arthropod limb. eLife 7:e34410. doi:10.7554/eLife.34410

Wolff T, Iyer NA, Rubin GM. 2015. Neuroarchitecture and neuroanatomy of the Drosophila central complex: A GAL4-based dissection of protocerebral bridge neurons and circuits. J Comp Neurol 523:997–1037. doi:10.1002/cne.23705

Yapici N, Cohn R, Schusterreiter C, Ruta V, Vosshall LB. 2016. A Taste Circuit that Regulates Ingestion by Integrating Food and Hunger Signals. Cell 165:715–729. doi:10.1016/j.cell.2016.02.061

Yu JY, Kanai MI, Demir E, Jefferis GSXE, Dickson BJ. 2010. Cellular Organization of the Neural Circuit that Drives Drosophila Courtship Behavior. Current Biology 20:1602–1614. doi:10.1016/j.cub.2010.08.025

Yu Y, Peng H. 2011. Automated high speed stitching of large 3D microscopic images2011 IEEE International Symposium on Biomedical Imaging: From Nano to Macro. Presented at the 2011 IEEE International Symposium on Biomedical Imaging: From Nano to Macro. pp. 238–241. doi:10.1109/ISBI.2011.5872396

Zhang YV, Ni J, Montell C. 2013. The molecular basis for attractive salt-taste coding in Drosophila. Science 340:1334–1338. doi:10.1126/science.1234133

Zheng Z, Lauritzen JS, Perlman E, Robinson CG, Nichols M, Milkie D, Torrens O, Price J, Fisher CB, Sharifi N, Calle-Schuler SA, Kmecova L, Ali IJ, Karsh B, Trautman ET, Bogovic JA, Hanslovsky P, Jefferis GSXE, Kazhdan M, Khairy K, Saalfeld S, Fetter RD, Bock DD. 2018. A Complete Electron Microscopy Volume of the Brain of Adult Drosophila melanogaster. Cell 174:730–743.e22. doi:10.1016/j.cell.2018.06.019

